# Selective Filopodia Adhesion Ensures Robust Cell Matching in the *Drosophila* Heart

**DOI:** 10.1101/222018

**Authors:** Shaobo Zhang, Christopher Amourda, Timothy E. Saunders

## Abstract

The ability to form specific cell-cell connections within complex cellular environments is critical for multicellular organisms. However, the underlying mechanisms of cell matching that instruct these connections remain elusive. Here, we explore the dynamic regulation of matching processes utilizing *Drosophila* cardiogenesis. During embryonic heart formation, cardioblasts (CBs) form precise contacts with their partners after long-range migration. We find that CB matching is highly robust at the boundaries between distinct CB subtypes. Filopodia in these CB subtypes have different binding affinities. We identify the adhesion molecules Fasciclin III (Fas3) and Ten-m as having complementary differential expression in CBs. Altering Fas3 expression influences the CB filopodia selective binding activities and CB matching. In contrast to single knockouts, loss of both Fas3 and Ten-m dramatically impairs CB alignment. We propose that differential expression of adhesion molecules mediates selective filopodia binding, and these molecules work in concert to instruct precise and robust cell matching.

## Introduction

Multicellular organisms contain a wide variety of differentiated cells that are specifically connected to their partners. Cell matching is the process that facilitates the formation of these connections within a complicated cellular environment that undergoes dramatic cell migration and rearrangement, and it is required for reliable embryogenesis and tissue remodeling (Dixon et al., 2011; Goodman and Shatz, 1993; Maragoudakis, 2000; Shinbane et al., 1997). Neurogenesis has been one of the favored models to understand how accurate cell matching occurs. Indeed, despite the complex cellular environment and dramatic cell rearrangement, neurons reliably recognize and form precise interconnections with their synaptic partners, and this is essential to the proper functions of the brain (Goodman and Shatz, 1993; Katz and Shatz, 1996; Woolf, 2000). Many other processes, such as neural crest formation (Sauka-Spengler and Bronner-Fraser, 2008), facial development (Dixon et al., 2011), angiogenesis (Adams and Alitalo, 2007) and wound healing (Martin, 1997), require similar cell matching to generate reliable cell-cell connections and eventually build robust biological architectures.

Filopodia, thin and actin-rich pioneer cell membrane protrusions, mediate cell matching through sensing the surrounding cellular environment (Davenport et al., 1993; Eilken and Adams, 2010; Jacinto et al., 2000; Tessier-Lavigne and Goodman, 1996). Recent studies have shown that filopodia in neuronal cells have selective stabilization properties and these dynamics specify growth cone stabilization and neural circuit formation (Hua and Smith, 2004; Özel et al., 2015; Xu et al., 2009). Various selective cell-cell adhesive molecules, such as Cadherins (Takeichi, 1987, 1988), immunoglobulin (Ig) superfamily proteins (Maness and Schachner, 2007; Williams and Barclay, 1988) and Teneurins (Hong et al., 2012; Mosca, 2015), have also been reported to instruct accurate cell matching, through homophilic and/or heterophilic interactions. Identification of these components involved in cell matching has largely been achieved through genetic studies in neurogenesis. However, their underlying mechanisms, especially the dynamic regulation of cell matching remains elusive, partially due to the complexity and limited accessibility for *in vivo* live imaging of the nervous systems. Here, we looked to investigate the process of cell matching in a simpler system - *Drosophila* embryonic heart formation - which facilitates quantitative live imaging of the cell matching process.

The *Drosophila* heart is a linear organ formed by two contralaterally symmetric rows of connected cardioblasts (CBs). These contralateral symmetric rows of CBs are initially around 100 μm apart. Concomitantly with dorsal closure, contralateral CBs collectively migrate towards the dorsal midline, closing the gap between the rows, and meet with their corresponding partners (Stage 16) (Bodmer and Frasch, 2010; Vogler and Bodmer, 2015). This process is reminiscent of the primitive heart tube formation in vertebrates (Bodmer, 1995; Srivastava and Olson, 2000). At the end of CB migration, heart cells from contralateral sides establish a one-to-one alignment (Figure 1A) and form a tube structure (Bodmer and Frasch, 2010; Vogler and Bodmer, 2015). Subsequently, the heart tube is divided into two domains: the anterior aorta and the posterior heart (Figure 1A). CBs are composed of distinct subtypes distributed in a repetitive fashion with four cells expressing the homeobox gene *tinman* (*tin*), and two expressing the orphan nuclear receptor gene *seven-up* (*svp*) (Figure 1A). This 4-2 cell arrangement persists throughout the CBs migration. In a fully formed heart, these CB subtypes give rise to distinct functional structures with significant morphological differences: Svp-positive CBs form ostia, while Tin-positive CBs constitute the major heart lumen and cardiac valves (Lehmacher et al., 2012; Medioni et al., 2009; Molina and Cripps, 2001). In vertebrate cardiomyopathy studies, cell misalignment can cause heart failure (Shinbane et al., 1997; Umana et al., 2003), indicating that proper contralateral cell matching is critical for heart function. Thus, as a confined system (continuous structure, predictable migration direction, and no cell division) amenable to *in vivo* live imaging and genetic perturbation (Bodmer and Frasch, 2010; Vogler and Bodmer, 2015), the formation of the *Drosophila* heart is an excellent system for studying the underlying mechanisms of cell matching.

**Figure 1.**
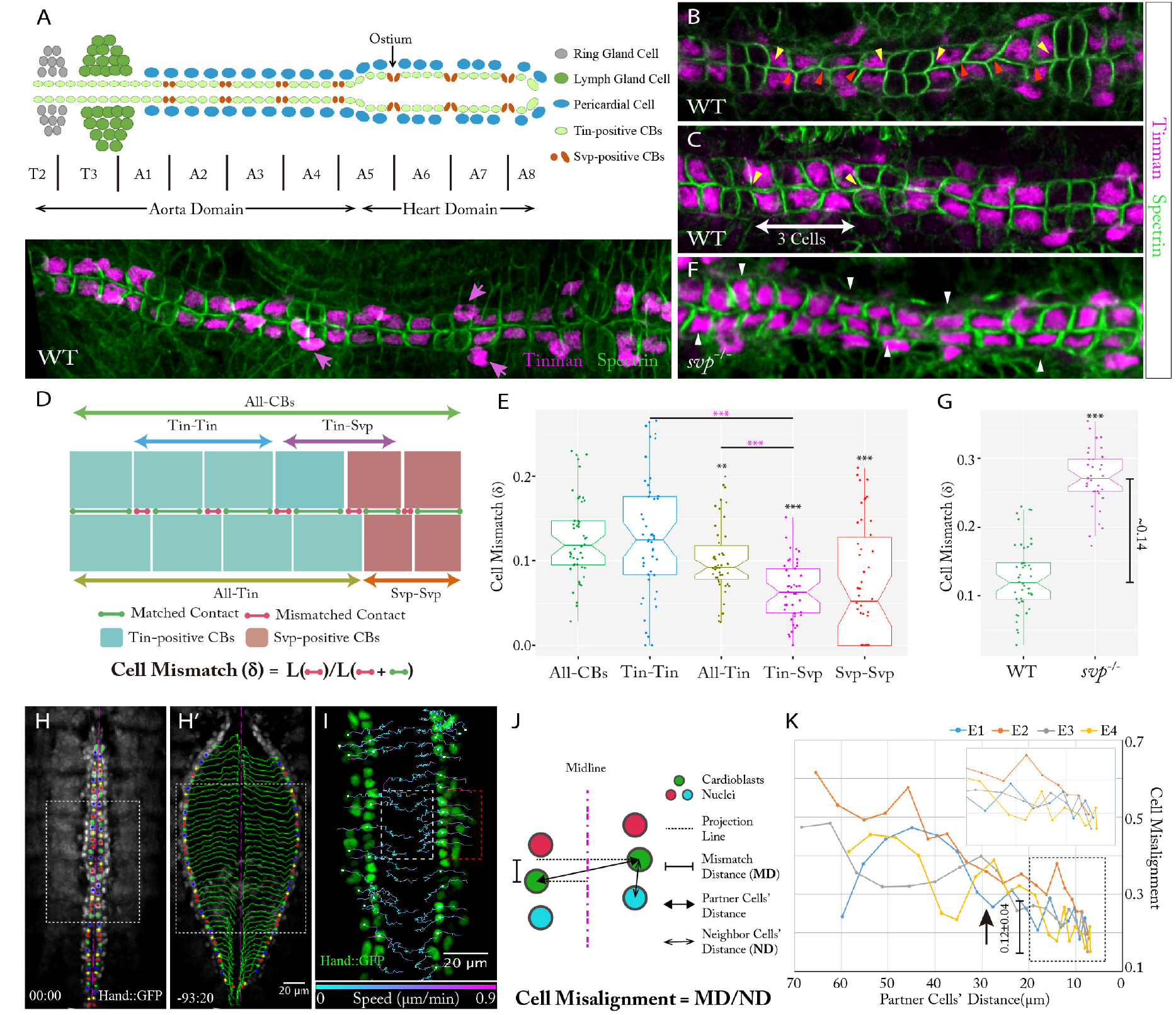
Active *Drosophila* CB matching is cell-type specific and short-ranged. (A) Top: Schematic representation of the *Drosophila* heart structure. Bottom: Immunostaining of the heart vessel with Tin and Spectrin at Stage 17. Pink arrows point to the Tin-positive pericardial cells. (B-C) CB membrane contacts in wildtype embryos at Stage 16. Yellow and red arrowheads point to examples of matched and mismatched membrane contacts. In (C) one of the cardiac segments has reduced Tin-positive CB number. (D) Schematic of cell mismatch definition (δ) between contralateral CBs. (E) Quantification of cell mismatch (δ) between different CBs in wildtype embryos (n=47). Data are presented as boxplot and scatter plot (*P<0.05, **P<0.01, ***P<0.001, ‘*’ - compared with ‘Overall’ contacts, ‘*’-compared with ‘Tin-Tin’ contacts). (F) Immunostaining of the heart vessel with Tin and Spectrin at Stage 17 in *svp*^−^ embryos. White arrowheads point to the cells with exaggerated mismatch to their contralateral neighbor. (G) Cell mismatch (δ) analysis of wildtype (n=47) and *svp*^−^ (n=34) embryos. Data are presented as boxplot and scatter plot (*P<0.05, **P<0.01, ***P<0.001). (H) Identification of contralateral cell partners (same color) at stage 16 in embryos expressing Hand::GFP. (H’) Cell tracking of same embryo as (H), with green lines showing the CB migration. (I) Individual CB cell tracking with shorter interval time (see also Figure S1E; Movie S2) before (red box) and after (white box) dorsal closure finishing with migration traces color-coded by migration speed. (J) Schematic of cell misalignment measurement during CB migration. (K) Quantification of the cell misalginment during CB migration (in region denoted by white boxes in H-J) (n=4). Inset is enlarged region highlighting cell matching at separations of <20 μm. Black arrow points to the rough time of dorsal closure finishing.

In this study, we show that active *Drosophila* CB matching primarily occurs at the boundaries between distinct CB subtypes. Hearts without different CB subtypes have dramatic CB mismatch. We reveal that CB filopodia have differential binding affinity in distinct cell types, with stronger adhesion between Tin-positive CBs. Through candidate screening, we identify that FasciclinIII (Fas3) and Ten-m (known as Tenascin-major in *Drosophila)* have complementary differential expression patterns in the heart. Fas3, belonging to the Ig superfamily (Chiba et al., 1995), shows higher expression in Tin-positive CBs. Changing Fas3 expression levels and pattern in CBs alters their filopodia binding activities and leads to CB mismatch. Ten-m, belonging to the Teneurins (Hong et al., 2012), shows higher expression in Svp-positive CBs. Losing Ten-m causes CB arrangement defects and cell mismatch. However, loss of either Ten-m or Fas3 does not fully abolish the active CB matching, as the other adhesion molecule appears able to partially compensate. In contrast, embryos without both Fas3 and Ten-m have dramatic CB matching defects, with a similar mismatch level to mutants without different CB subtypes. Therefore, our results suggest that differential adhesion regulates the filopodia selective binding activity and this provides a simple but efficient mechanism to instruct precise and robust cell matching.

## Results

### Active *Drosophila* CB matching is cell-type dependent and short ranged

After long-range migration, CBs typically form near-perfect alignment with their partners from the contralateral side (Figure 1A) (Bodmer and Frasch, 2010; Vogler and Bodmer, 2015). Yet, malformed hearts are observed in ~8% (7/84) of wild-type embryos, potentially due to natural variation. To test whether cell alignment is required for heart function, we imaged the heart beating in embryos where CBs are labeled with Hand::GFP (Han, 2006). Compared to well-matched hearts (Figure S1A and Movie S1 Left), both the morphology and function are impaired in hearts with misaligned CBs (Figure S1B and Movie S1 Right). In these embryos with misaligned CBs, the nuclear separation between contralateral CB partners in the aorta domain reduces from ~6 to ~4 μm and the heart beating does not propagate through the mismatched regions.

Through closer examination of the membrane contacts between contralateral CBs in wildtype embryos, we noticed that contact mismatch between CBs of different subtypes is quite rare, but relatively common between CBs of the same subtype (Figure 1B). Even in embryos where the 4-2 pattern of the Tin- and Svp-positive CBs is disrupted (n=15/84), the cell matching remains precise at the CB subtype boundaries (Figure 1C). Quantifying the CB contact mismatch (denoted by δ) in the aorta region (Figure 1D), we found that contacts of adjacent Tin- and Svp-positive CBs show considerably smaller degree of mismatch when compared with the adjacent Tin-positive CBs (δ_Tin-Svp_=0.06±0.03, δ_Tin-Tin_=0.13±0.07, Figure 1E). To test whether distinct cell types are necessary for precise CB alignment, we investigated the cell matching in *svp*^−^ mutant embryos, where all the CBs are Tin-positive (Lo and Frasch, 2001). We found that such hearts show severe matching defects (Figure 1F) with dramatically higher δ compared with wild-type embryos (δ_WT_=0.13±0.05, δ_*svp−*_=0.27±0.04, Figure 1G). Moreover, in embryos with the *TM3* balancer, which carries a mutation in the CB patterning gene *Ubx* (Ponzielli *et al*., 2002), we found that the distinct CB subtypes are able to align, except for the regions with severely perturbed CB arrangement (Figures S1C and S1D). These results suggest distinct cell types are necessary for robust *Drosophila* CB matching.

To further explore when and where active CB matching happens, we tracked Hand::GFP labeled CBs throughout heart closure (Figure 1H-1I) and measured the misalignment between partner CBs based on their centroids (Figures 1J and 1K, Methods). The CB misalignment gradually decreases throughout formation of the heart (Figure 1K). However, once CBs approach within a separation of ~15-20 μm, after the completion of dorsal closure (Figure S1E), positional readjustments relative to their contralateral partners are more frequent (Figure 1I and 1J, Movie S2), and correspondingly rapid changes in the cell misalignment quantification are observed (Figure 1K). Interestingly, the increase in cell alignment during this active stage of CB positional adjustment (0.12±0.04, Figure 1K) is comparable to the difference in the measured contact mismatch (δ) between *svp−* and wildtype embryos (δ_*svp*^−^_ - δ_WT_=0.14, Figure 1G). Further, by imaging Hand::GFP labeled CBs throughout cardiogenesis, we observed no significant changes in the relative position of CBs after the contralateral sides fully meet at the midline (Figure S1F).

Taken together, active CB matching requires proper heart cell differentiation and occurs when the contralateral cells are within ~15-20 μm apart, which is after dorsal closure and precedes the complete coalescence of the contralateral rows of CBs.

### Filopodia show differential binding affinity in distinct CB subtypes

We next sought to identify the subcellular components that are responsible for accurate CB matching. Considering the active CB matching happens within a short distance, we explored filopodia activity during the matching process. Filopodia are known to guide cell recognition and short range targeting in other systems (Davenport et al., 1993; Eilken and Adams, 2010; Jacinto et al., 2000; Özel et al., 2015). To test whether they also regulate heart cell matching, we imaged the CB filopodia activity (Figure 2A) by driving Moesin (Moe)::GFP (Edwards et al., 1997) expression in all CBs using Hand-Gal4 (Han, 2006). As reported previously (Haack et al., 2014; Swope et al., 2014), CBs filopodia are highly dynamic, protruding and retracting repeatedly (Movie S3 Left). At a cell center separation of ~15-20 μm, we observed that contralateral CBs initiate physical contacts through filopodia (Figure 2A). This process persists for around 30 minutes until the two sides completely adhere with each other. Closer examination of the matching process showed that after forming filopodia contacts, CBs adjust the relative matching position towards their contralateral partners (Figure 2A).

**Figure 2.**
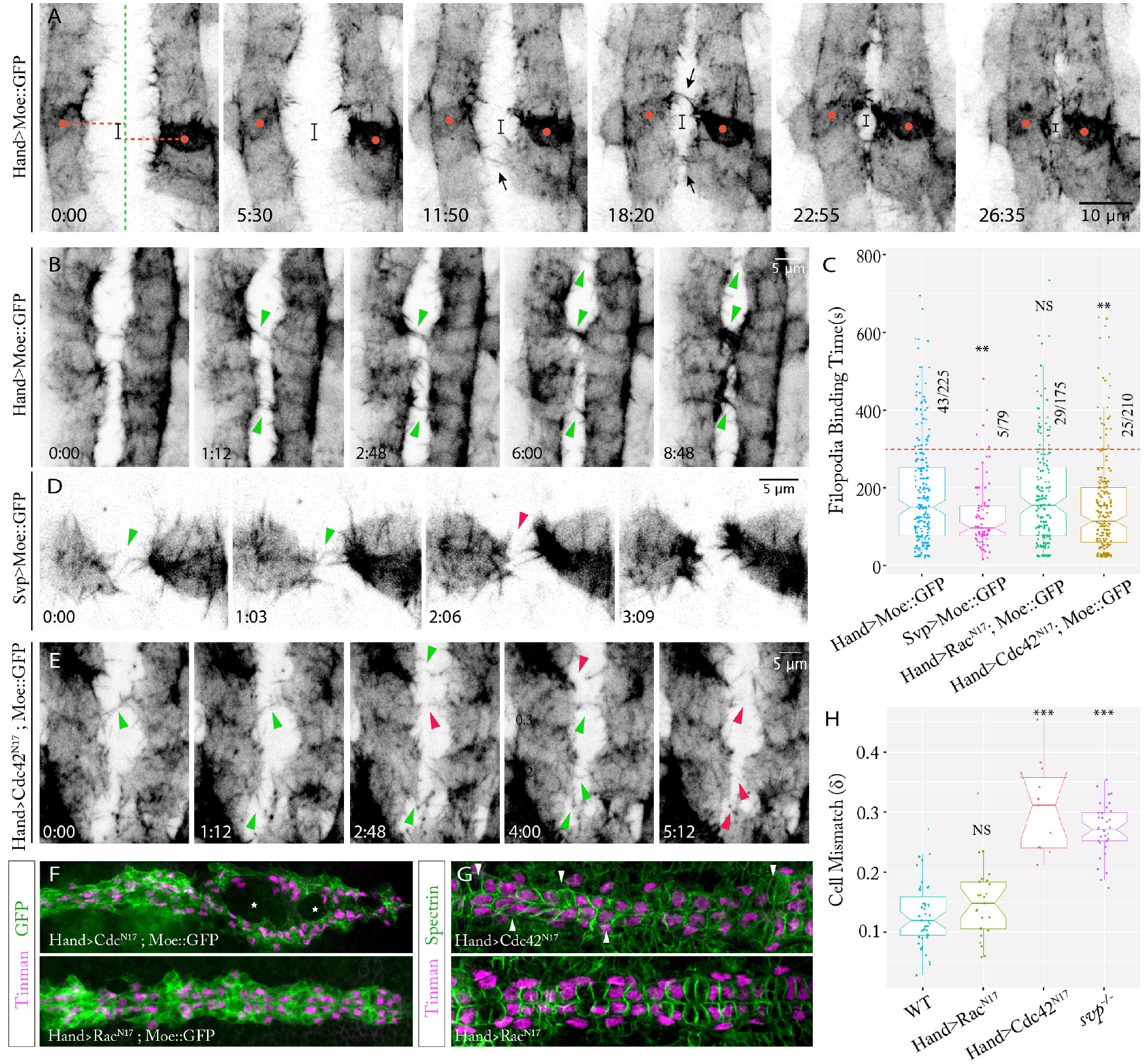
CB filopodia drive cell matching through differential binding affinity in distinct CB subtypes. (A-B) CB filopodia activity in Hand>Moe::GFP embryos. In (A) black bar denotes contralateral partner cell (denoted by red dots) along the midline (green line) offset during migration (see also Movie S3 Left). (C) Quantification of filopodia binding time under different conditions. Data are presented as boxplot and scatter plot (n>70 per condition) (*P<0.05, **P<0.01, ***P<0.001, NS: Not Significant, compared with Hand>Moe::GFP condition), red line corresponds to 300s binding time. (D) Filopodia activity in a Svp>Moe::GFP embryo (see also Movie S4 Left). (E) Filopodia binding activity in Hand>Cdc42^N17^; Moe::GFP embryo (see also Movie S3 Right). (B, D, E) Green and red arrowheads point to reinforced and transient filopodia adhesions respectively. (F) Heart morphology in Hand>Cdc42^N17^ (top) and Hand>Rac^N17^ (bottom) embryos (both expressing Moe::GFP) at stage 16. Asterisks label unclosed gaps between contralateral sides. (G) CB alignment in Hand>Cdc42^N17^ (top) and Hand>Rac^N17^ (bottom) embryos. White arrowheads point to the cells with exaggerated mismatch. (H) CB mismatch (δ) quantification in different conditions. Data are presented as boxplot and scatter plot (n>5 per condition) (*P<0.05, **P<0.01, ***P<0.001, NS: Not Significant, compared with WT).

Critically, we found that CB filopodia have selective binding properties. After initial contact, a subset of filopodia from opposing cells stabilize their contacts and show accumulated Moe::GFP signal (Figure 2B and Movie S3 Left), indicating increased actin-related activities. Eventually, these stabilized filopodia enlarge their contact area and direct the corresponding CBs toward each other. We found that the binding time of filopodia contacts for all CBs is around 185±141s. This large variation is explained by the fact that 19% of observed contacts persisted over 300s whilst 33% lasted less than 100s (Figure 2C). In embryos where distinct CB subtypes can be distinguished (Figure S2A-S2B), strong filopodia binding is observed forming between Tin-positive CBs (Figure S2B). To further determine whether this selective binding property is related to distinct cell types, we examined the filopodia activity in different CB subtypes using cell specific Gal4 drivers. When driving Moe::GFP expression under TinC-Gal4 (Lo and Frasch, 2001), we observed leaking expression of Moe::GFP, not only in Svp-positive CBs but also in the aminoserosa cells between the contralateral CBs (Figure S2C). In contrast, Moe::GFP driven by Svp-Gal4 (Pfeiffer et al., 2008) shows specific expression (Figure 2D). In these embryos, we found that the filopodia contacts between Svp-positive CBs rarely stabilize (Figure 2D and Movie S4 Left), and show much shorter binding time (averaged binding time=130s, with 6.3% of contacts persisting over 300s, Figure 2C). Adhesion between these Svp-positive CBs does not occur until larger lamellipodia contacts form (Movie S4 Left). These Svp-positive CBs are also more rounded during migration, which appears to minimize their contact with the neighboring cellular environment. Altogether, these data suggest that filopodia of Tin-positive CBs have higher binding affinity compared with the ones of Svp-positive CBs.

As the molecular mechanisms of CB filopodia formation and activity are still unknown (Haack et al., 2014), we perturbed filopodia activity in CBs by expressing a dominant-negative allele of Cdc42 (Cdc42^N17^) (Luo et al., 1994) using Hand-Gal4. Cdc42 is a small GTPase that regulates polarity establishment, cell migration and filopodia activity (Hall, 1998; Mattila and Lappalainen, 2008), and in contrast to Rac and Rho, it is essential for heart morphogenesis and lumen formation (Swope et al., 2014; Vogler et al., 2014). Expressing Cdc42^N17^ in the heart, we observed no significant change in the density of CB filopodia compare with control (0.45±0.15 filopodia/μm in Cdc42^N17^, 0.47 ±0.07 filopodia/μm in control, Figure S2A). However, Cdc42^N17^ expression leads to decreased filopodia binding affinity and loss of obvious Moe::GFP accumulation (Figure 2E and Movie S3 Right), with corresponding reduction in filopodia binding time (averaged binding time=150.9s, with 11.9% persisting more than 300s, Figure 2C). Heart morphology in Cdc42^N17^ expressing embryos is dramatically perturbed, with many unclosed gaps between contralateral CBs (18/34 embryos), CB misalignment and high cell mismatch (δ=0.31±0.07, Figure 2F-2H). In contrast, when expressing a dominant-negative allele of the small GTPase Rac (Rac^N17^) (Luo et al., 1994) under the same Hand-Gal4 driver, we observed no obvious changes in either filopodia density (0.46±0.10 filopodia/μm, Figure S2D) or filopodia binding activity (averaged binding time=179.2s, with 16.6% persisting more than 300s, Figure 2C and Figure S2E). Examining the heart morphology, we did not observe either significant perturbation of heart structure or CB alignment in Rac^N17^ expressing embryos (δ=0.15±0.06, Figure 2F-2H). Taken together, we propose that the selective filopodia adhesion in distinct CB subtypes plays a critical role in regulating CB matching.

### Candidate screening identified differential expression of Fas3 and Ten-m in the heart

To identify the specific molecules that directly mediate the differential CB filopodia adhesion and cell matching, we performed a candidate-based screen. Candidate molecules (Figure S3A) were selected using the following criteria: located at the cell surface; known to regulate cell-cell adhesion; and reported to mediate cell recognition and matching in other systems. We examined the expression of these candidates in the *Drosophila* heart - through immunostaining - to identify the ones showing differential expression in distinct CB subtypes. Neuroglian, Connectin, Fasciclin I and Fasciclin II have no detectable expression in the heart; Cadherins and Neurotactin are uniformly distributed throughout the CBs (Figure S3B-S3B”). In contrast, Fas3 and Ten-m show noticeable differential expression patterns within the *Drosophila* heart (Figures S3C and S3C’). Both Fas3 (Chiba et al., 1995; Kose et al., 1997; Snow et al., 1989) and Ten-m (Hong et al., 2012; Mosca, 2015) are reported to regulate synaptic target matching through promoting homophilic attraction. Further co-staining with Fas3 and Tin antibodies, we found that Fas3 is highly expressed in Tin-positive CBs (Figure 3A), while at a much lower level in Svp-positive CBs. Confirmed through co-staining in both wild-type and endogenous Ten-m::GFP tagged embryos (MiMIC-Ten-m::GFP) (Nagarkar-Jaiswal et al., 2015), Ten-m expression shows a complementary expression pattern to Fas3, with relatively higher expression in Svp-positive CBs than Tin-positive CBs (Figure 3B and Figures S3D and S3D’).

**Figure 3.**
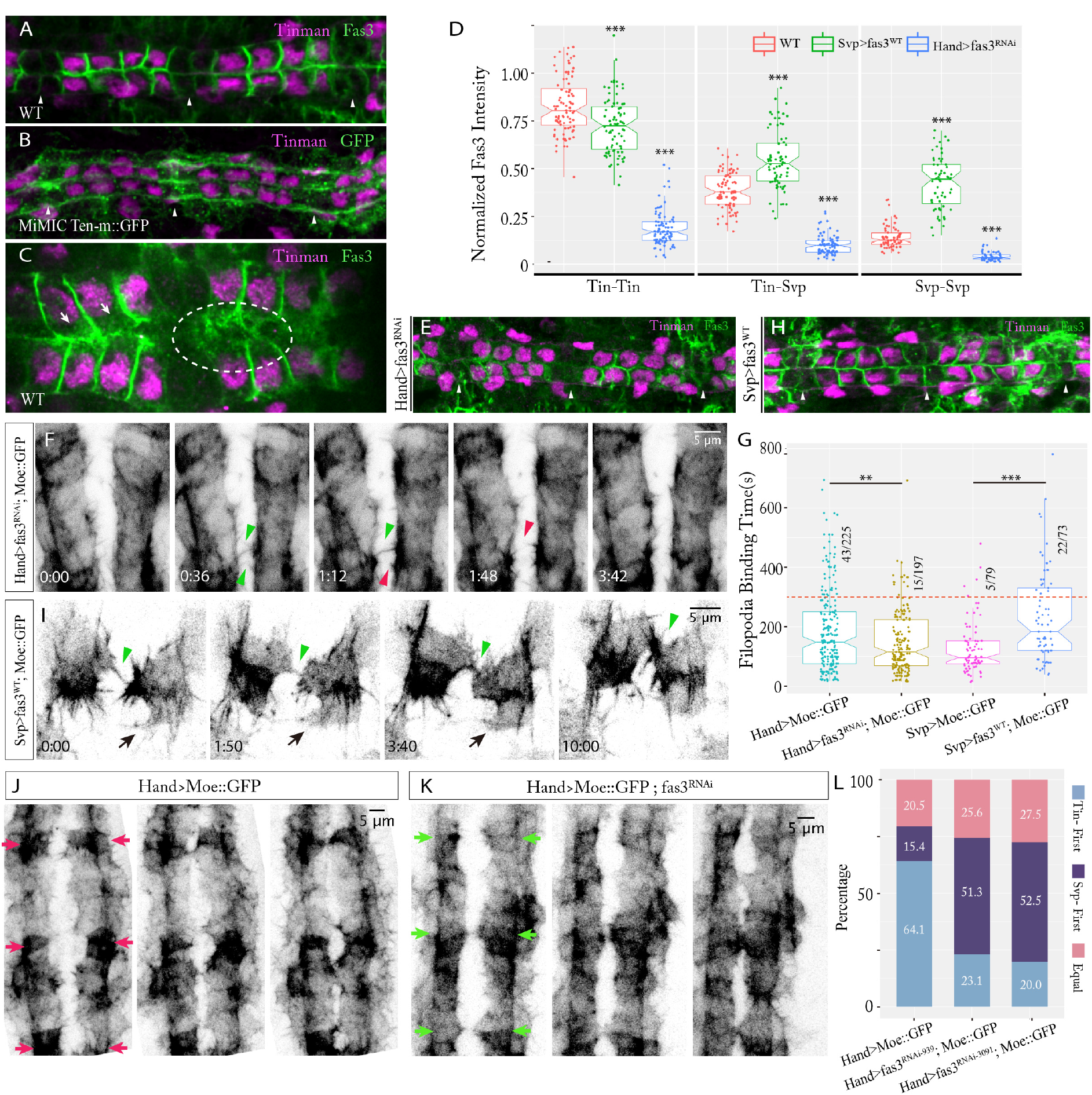
Fas3 shows differential expression in *Drosophila* heart and modulates CB filopodia adhesion. (A, B) Expression patterns of Fas3 (A) and Ten-m (B) in the heart at early Stage 17. (C) Fas3 localization between contralateral CB partners at Stage 16, highlighted by dashed ellipse. Arrows point to Fas3 pools near the forming cell-cell junctions. (D) Normalized Fas3 intensity at different cell-cell contacts under different experimental conditions (n=7 embryos per condition). Data are presented as boxplot and scatter plot (*P<0.05, **P<0.01, ***P<0.001 compared to the wildtype at each different cell-cell contact). (E) Hand>fas3^RNAi^ embryos showing reduced Fas3 expression in CBs. (F) Filopodia activity in Hand>fas3^RNAi^ embryos expressing Moe::GFP embryo (see also Figure S4C-S4D; Movie S3 Middle). (G) Quantification of filopodia binding time under different Fas3 expression conditions (see also Figure S4E). Data are presented as boxplot and scatter plot, (*P<0.05, **P<0.01, ***P<0.001). (H) Fas3 overexpression in Svp-positive CBs. (A, B, E, H) Arrowheads point to Svp-positive CBs. (I) Filopodia activity in Svp>fas3^WT^embryos expressing Moe::GFP (see also Movie S4 Right). (F, I) Green and red arrowheads point to reinforced and transient filopodia adhesions respectively. (J, K) CB migration pattern in embryos expressing Moe::GFP under control (J) and Hand>fas3^RNAi^ (K) conditions. Green and red arrows highlight Svp-positive CBs that initiate membrane contact and fuse earlier or later than Tin-positive cells respectively (see also Movie S5). (L) Percentage of different migration patterns under different experimental conditions (n>30 per condition); Tin-positive CBs initiate contact (blue), Svp-positive CBs initiate contact (purple), Tin- and Svp-positive CBs initiate contact at equal time (red).

### Differential Fas3 expression level modulates filopodia adhesion in CBs

The Fas3 expression pattern in the heart coincides with the differential CB filopodia binding affinity. We next examined more closely Fas3 expression and function. Fas3 is observed at the newly formed cell-cell junctions (Figure 3C), especially between Tin-positive CBs. Pools of Fas3 are found near the forming junctions, indicating potential active transport of Fas3 towards the cell leading edge to stabilize these contacts. Further, Fas3 localizes to the regions where filopodia contacts between contralateral matching CBs form (Figure 3C). Through exploring Fas3 expression during CB migration, we find that Fas3 expression is not only spatially but also temporally regulated during cardiogenesis (Figure S3E-S3H’). At Stage 14, there is no obviously detectable Fas3 expression, but as the cells migrate toward the embryo midline, Fas3 expression emerges and gets steadily higher with a differential expression pattern noticeable. In tin^ABD^; tin^346^/tin^346^ embryos, where Tin expression is depleted in the CBs during their migration (Zaffran, 2006), the differential expression of Fas3 is lost and the CB matching is dramatically affected (Figures S3I and S3I’).

To test the role of Fas3 in CB matching - particularly in regulating selective filopodia binding activity - we performed *fas3* RNA interference (RNAi) in all CBs using Hand-Gal4, and Fas3 overexpression in Svp-positive CBs using Svp-Gal4. Both *fas3* RNAi lines used (fas3^RNAi-939^ and fas3^RNAi-3091^) efficiently decrease Fas3 expression in CBs (Figures 3D and 3E and Figure S4A). When Fas3 expression dramatically declines, the majority of CB filopodia contacts fail to stabilize (Figure 3F and Figures S4B and S4C and Movie S3 Middle), and show noticeably shorter filopodia binding time (averaged binding time=145s, with 8% persisting over 300s using fas3^RNAi-3091^; averaged binding time =150s with 15% persisting over 300s using fas3^RNAi-939^, Figure 3G and Figure S4D). In contrast, UAS-fas3^WT^ driven by Svp-Gal4 markedly increases the Fas3 expression in Svp-positive cells (Figures 3D and 3H). Filopodia in the Svp-positive CBs with upregulated Fas3 expression show increased binding affinity, with contacts stabilizing and Moe::GFP accumulating (Figure 3I and Movie S4 Right). In these cells, the filopodia binding duration significantly increases (averaged binding time = 230s, with 30% persisting over 300s, Figure 3G). Further, the morphology of Svp-positive CBs is drastically changed, with significant cell expansion and large, long-lived protrusions, indicative of increased adhesion to neighboring cells (Figure 3I). These results reveal that the spatially varied Fas3 expression level is critical for differential filopodia adhesion between CBs.

We further noticed that decreasing the Fas3 expression level in CBs also changes their collective migratory behavior. CBs join with their contralateral partners through a ‘buttoning’ pattern, with Tin-positive CBs typically making initial contact (Figures 3J and 3L and Movie S5 Left) (Swope et al., 2014). However, when Fas3 expression is reduced, the sequence of heart closure is reversed, with Svp-positive CBs contacting first with their contralateral partners (Figures 3K and 3L and Movie S5 Right). This suggests that Fas3 not only regulates filopodia binding activity but also the collective CB migration. As the epidermis above the CBs shows strong Fas3 expression (Figure S4E and S4E’), Fas3 may also be acting to guide CBs migration on the epidermal sheets, especially for Tin-positive CBs that have high Fas3 expression. However, Fas3 is less critical in the migration of Svp-positive CBs, as their migration is less perturbed by reduction in Fas3 expression, suggesting that other molecules (potentially Ten-m) mediate the Svp-positive CB migration and fusion.

### Disruption of differential Fas3 expression pattern alters CB matching

The Fas3 expression level determines the corresponding CB filopodia binding affinity. Therefore, its differential expression pattern in the heart should be critical for CB matching. To test this hypothesis, we generated various Fas3 expression patterns in the developing *Drosophila* heart using: *fas3* null mutant (Fas3^E25^) (Elkins et al., 1990); cell-specific *fas3* RNAi; and Fas3 overexpression (Figure 4A-4F). Down-regulating expression of Fas3 in Svp-positive CBs - thereby increasing the relative Fas3 expression difference between Tin- and Svp-positive CBs - does not significantly alter cell matching, with the cell boundaries remaining well-matched in the aorta domain (δ=0.13±0.04, Figures 4B-4B”’ and 4G). In Fas3^E25^ embryos, we observed mismatched CB contacts more frequently (Figure 4C-4C”’), with a significant, though relatively small, increase in the cell mismatch (δ=0.19±0.11, Figure 4G). We also noticed that in these embryos the CBs shape becomes more rounded, a phenotype reminiscent of Svp-positive CBs (Figures 4C” and 4C”’). Next, we tested the reverse condition, where all the CBs have approximately equal level of Fas3, by increasing Fas3 expression in Svp-positive CBs (Figure 4D-4D”’). We found a slightly increased level of CB matching defects compared to Fas3^E25^ embryos (δ=0.22±0.06, Figures 4D”, 4D”’ and 4G). Noticeably, Fas3 overexpression leads to more angular (square-like) cell shapes in Svp-positive CBs (Figures 4D” and 4D”’). These Fas3 perturbation results confirm that the differential expression pattern of Fas3 is important for CB shape and precise matching.

**Figure 4.**
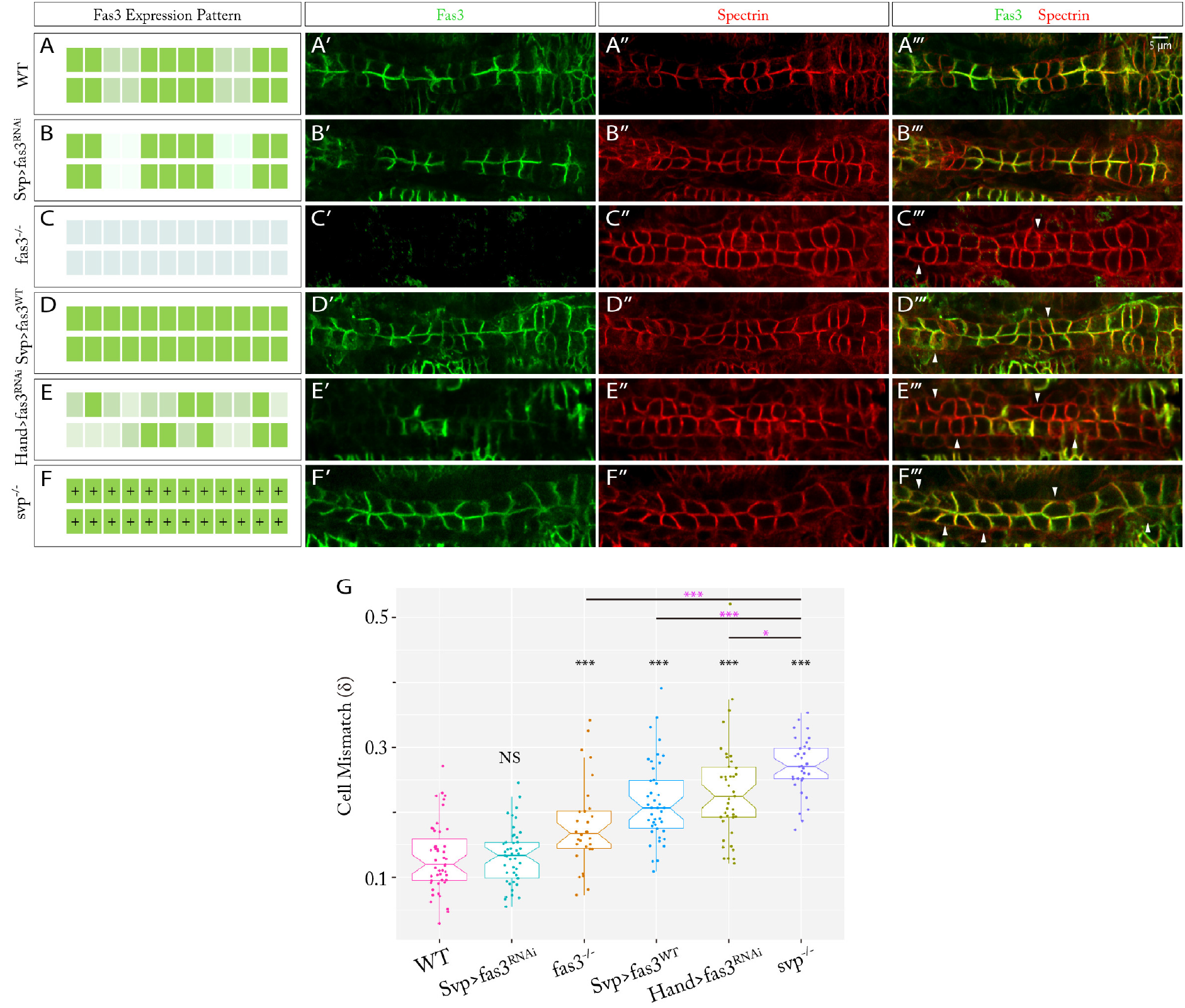
Disruption of the Fas3 expression pattern alters CB matching. (A-F) Schematics of various Fas3 expression patterns tested in the heart. Shades of green indicate the various Fas3 expression levels, ‘+’ indicates all CBs are same cell type. (A’-F’) Fas3 staining expression patterns under different conditions. (A”-F”) CB membrane contacts shown by Spectrin staining. (A”’-F”’) Merged images of Fas3 and Spectrin staining. Arrowheads highlight clear cell mismatch. (G) Cell Mismatch (δ) analysis of different Fas3 expression conditions (n>20 per condition). Data are presented as boxplot and scatter plot (*P<0.05, **P<0.01, ***P<0.001, NS: Not Significant, compared with WT).

To further challenge the matching capacity of CBs, we generated stochastic Fas3 expression patterns by taking advantage of the large cell-to-cell variability in activity of the Hand-Gal4 driver (Figure S5A-S5C; see Supplementary Information for Hand-Gal4 activity). Crossing Hand-Gal4 with fas3^RNAi^ created hearts with significant Fas3 intensity variation (Figure 4E-4E”’ and Figures S5D and S5E). Under this condition, CB matching is dramatically perturbed and shows even larger mismatch than Fas3^E25^ embryos (δ_RNAi-3091_=0.23 ±0.08, δ_RNAi-939_=0.22±0.07, Figures 4E”-4E”’ and 4G and Figure S5F-S5H”). Moreover, we observed that within the same embryo, the subset of CBs with higher Fas3 expression tended to be larger and maintained a more angular shape, compared with cells having low Fas3 expression (Figure 4E”’and Figures S5G” and S5H”). This result gives further evidence of the important role of Fas3 for CB morphology and matching. However, despite the dramatically perturbed CB alignment in the above Fas3 perturbations, the highest cell mismatch is still observed in *svp*^−^ embryos (Figure 4F-4G), which lose all the cell-type based matching mechanisms, indicating compensated matching mechanisms potentially exist.

### Low expression of Fas3 in Svp-positive CBs is necessary for robust heart morphogenesis

Low, instead of null, Fas3 expression exists in Svp-positive CBs (Figure 5A-B”). To test whether low Fas3 expression level plays a role in cardiogenesis, we examined the Svp-positive CB matching in Svp>fas3^RNAi^ embryos, where Fas3 expression in Svp-positive CBs is noticeably reduced (Figures 5C and 5C’). We observed a markedly increased Svp-positive CB contact defects in the heart domain (Figure 5A), where the Svp-positive CBs fail to make any contact with their partners (Figure 5C”) in more than 20% of Svp>fas3^RNAi^ embryos (17/74 using fas3^RNAi-3091^ and 11/40 using fas3^RNAi-939^), compared with wildtype embryos (8/84, p=0.02). Next, we tested the reverse condition by overexpressing Fas3 in the heart using UAS-fas3^WT^ and Hand-Gal4 which results in particularly increased Fas3 expression in Tin-positive CBs (Figures S6A and S6A’). When this over-expression was especially marked in the heart domain, we observed similar Svp-positive CB contact defects (Figure D-D”), but with even higher frequency (24/78). In contrast, overexpression of Fas3 specifically in Svp-positive CBs shows comparable defect frequency (8/78) to wild-type embryos. In the heart domain, contacts between contralateral Svp-positive CBs are typically narrower (contralateral contact length=1.7±0.5μm) than the ones in the aorta domain (contralateral contact length=3.2±0.4μm) (Figure S6B). As the filopodia in CBs can reach up to ~5μm, the diagonally opposite Tin-positive CBs neighboring Svp-positive CBs are able to make crossover contacts in the heart domain (Figures 5E and 5F and Movie S6). In normal condition, these crossover contacts rarely stabilize (Figure 5F and Movie S6). Hence, we propose that the basal Fas3 expression in Svp-positive CBs may act to inhibit the stabilization of these crossover contacts through mediating filopodia binding competition (Figures 5E) and therefore ensure robust heart morphogenesis. Combined, these results suggest that the low expression of Fas3 in Svp-positive CBs is necessary and the expression difference between Tin-positive CBs and Svp-positive CBs needs to be precisely regulated.

**Figure 5.**
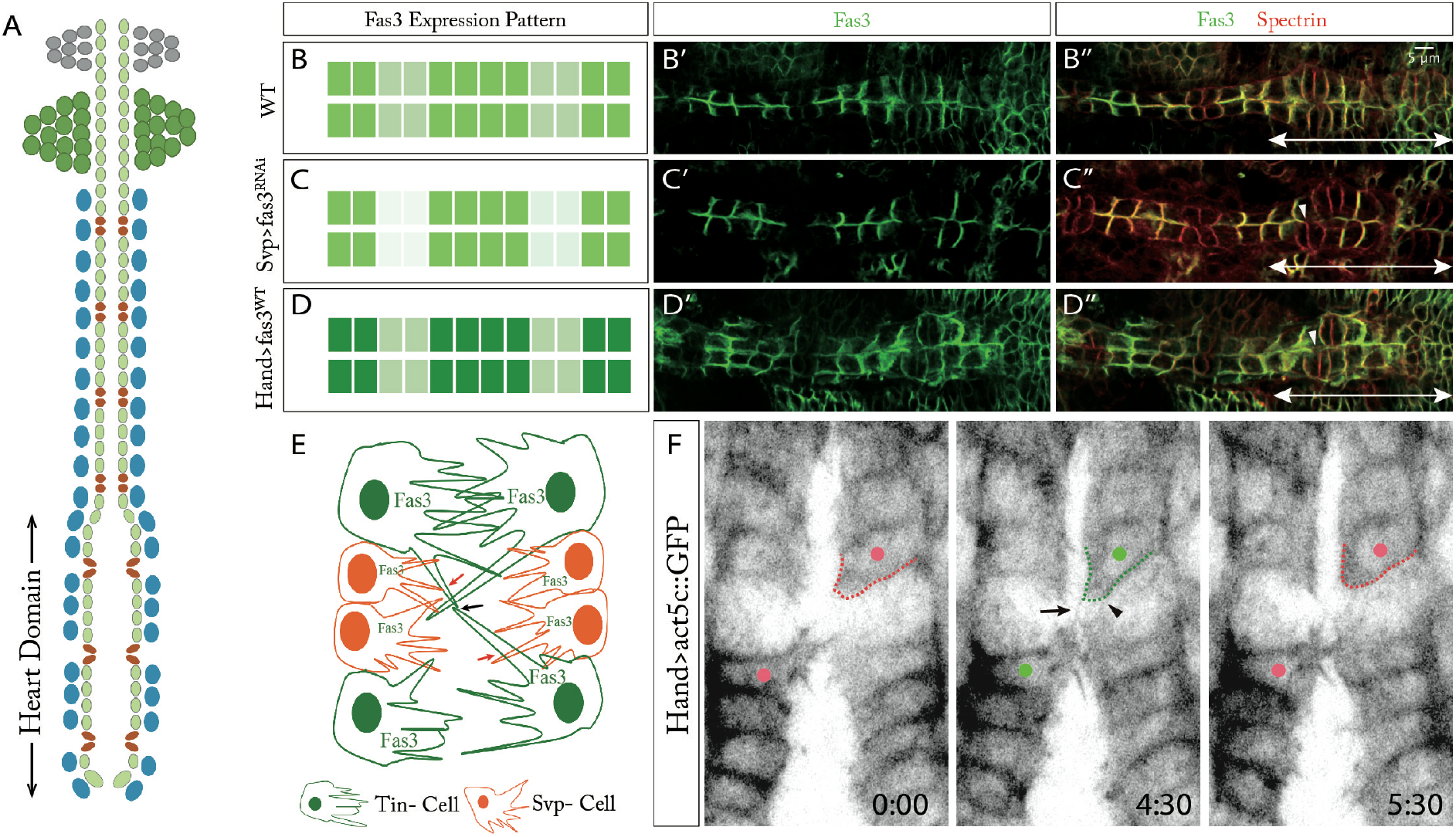
Low expression of Fas3 in Svp-positive CBs is necessary for robust heart morphogenesis, especially in the heart domain. (A) Schematic representation of the *Drosophila* heart structure, heart domain is labeled out. (B-D) Schematics of Fas3 expression patterns tested in the heart in of wildtype, Svp>fas3^RNAi^ (H, H’) and Hand>fas3^WT^. (B’-D’) Fas3 staining in the heart under different conditions. (B”-D”) Merged images of Fas3 and Spectrin staining under different conditions. Arrowhead highlights severe mismatch between Svp-positive CBs. Double-headed arrows show the heart domain. (E) Schematic demonstration of Fas3 function in Svp-positive CBs. Black arrows point to the crossover contacts between diagonal Tin-positive CBs that are neighboring Svp-positive CBs. Red arrows point to the filopodia contacts between Svp- and Tin-positive CBs. (F) Crossover filopodia contacts (black arrow pointed) between diagonal Tin-positive CBs in Hand >act5c::GFP embryos (see also Movie S6). Green and red dots label the Tin-positive CBs with or without crossover contacts, green and red dash lines label the Tin-positive CBs shape change with or without crossover contact.

### Ten-m works complementarily with Fas3 to ensure robust CB matching

Our above results suggest redundancy or complementary matching mechanisms to Fas3 exist to ensure robust cell matching in the heart. As shown through our candidate screening, Ten-m has complementary differential expression to Fas3 in *Drosophila* heart, with higher expression in Svp-positive CBs (Figure 3B). We next examine the expression and functional relationships between Ten-m and Fas3 in detail. Fas3 staining in MiMIC Ten-m::GFP embryos, confirmed the reversed expression pattern of Ten-m and Fas3 in the heart (Figure 6A-6B”). At the interface between the contralateral CBs, we found that Ten-m and Fas3 are expressed at high levels in distinct domains, with few exceptions showing either both high or low levels of both Ten-m and Fas3 (Figure 6C).

**Figure 6.**
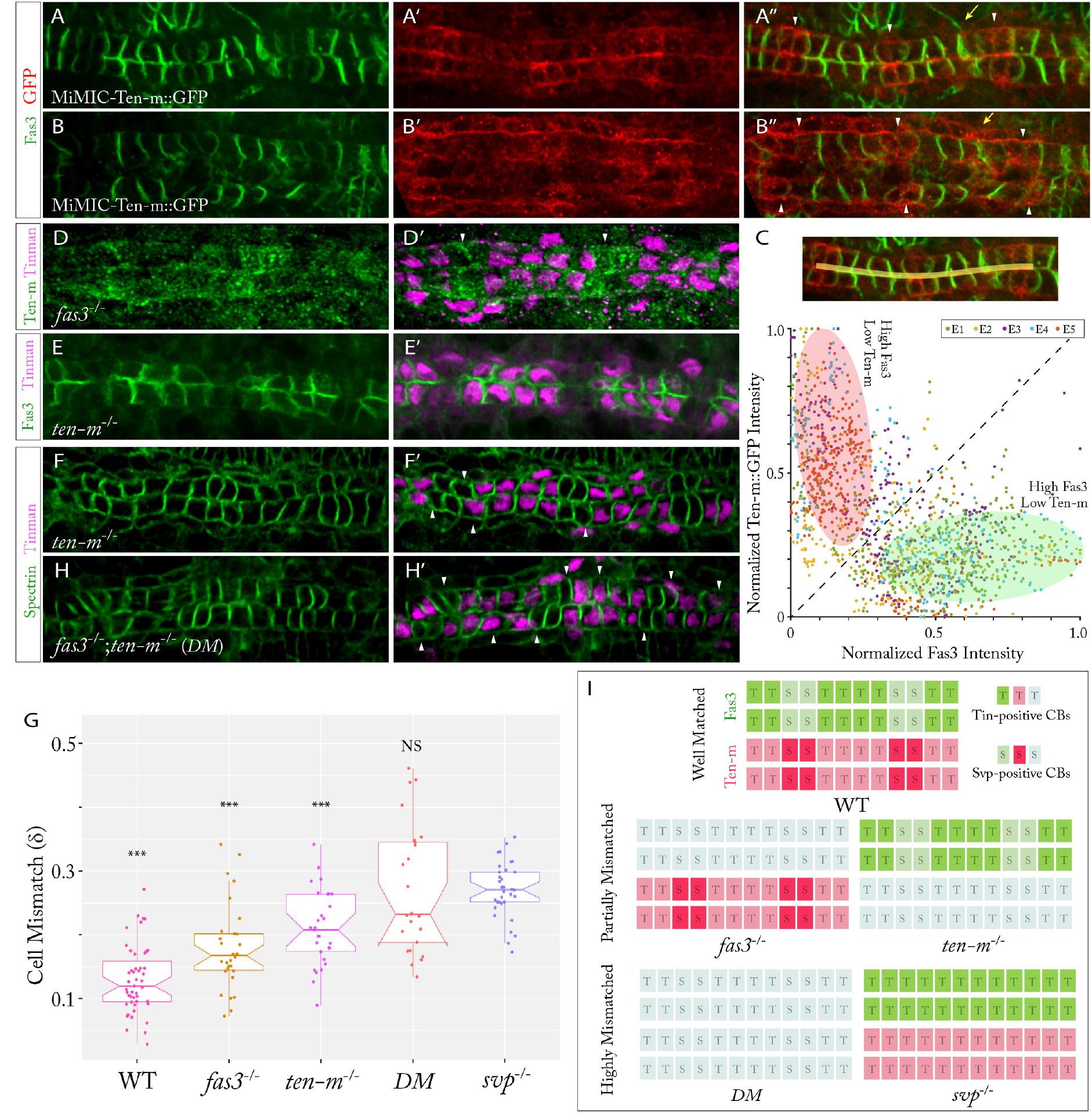
Ten-m and Fas3 regulate CB matching in a complementary fashion. (A-B”) Fas3 and GFP staining in MiMIC Ten-m::GFP embryos at Stage 17 (A) and Stage 16 (B). Arrowheads highlight regions of higher Ten-m expression, yellow arrows point to the pericardial cells. (C) Comparison of Fas3 and Ten-m relative intensity at contralateral CB contacts (yellow line in inset) (n=5). Green shade labels the region with high Fas3 expression but low Ten-m, red shade labels the reversed pattern. (D-D’) Ten-m staining expression in fas3- embryo. Arrowheads highlight higher Ten-m expression. (E-E’) Fas3 and Tin staining in ten-m- embryo. (F-F’) CB alignment in ten-m- embryo stained for Tin and spectrin. (G) Cell mismatch (δ) quantification of different experimental conditions (n>20 per condition). Data are presented as boxplot and scatter plot (*P<0.05, **P<0.01, ***P<0.001, NS: Not Significant, compared with svp-). (H-H’) CB alignment in fas3-; ten-m- embryo stained for Tin and Spectrin. Arrowheads highlight clear cell mismatch. (I) Schematics of Ten-m and Fas3 complementary functions in CB matching.

In contrast to Fas3, Ten-m is not only found in the CBs but also in the surrounding pericardial cells (Figures 6A” and 6B”) and shows much earlier expression in CBs (Figure S6A-S6B”’), with transcriptional activity reported at Stage 12 (Baumgartner et al., 1994). Compared to Fas3, Ten-m expression in the epidermis is more localized, and, strikingly, these regions correspond to the locations above Svp-positive CBs (Figure S6C-S6C’). Potentially, Ten-m plays a role in mediating Svp-positive CBs migration through providing an adhesive environment, especially in embryos with reduced levels of Fas3 in the CBs. This gives an explanation of the reversal in the collective migration pattern of CBs observed in Hand>fas3^RNAi^ embryos (Figure 3K).

If Ten-m and Fas3 are acting as complementary mechanisms to ensure robust CB morphogenesis, their expression patterns should be independent from each other within CBs; *i.e*. losing one does not affect the differential expression of the other. To test this, we checked Ten-m expression in *fas3^-^* mutants and Fas3 expression in Ten-m null embryos (Ten-m [5309]) (Levine et al., 1994). As expected, loss of one factor does not change the other’s expression pattern (Figure 6D-6E’).

To test whether Ten-m and Fas3 are working in concert to ensure robust CB matching, we first examined the cell arrangement and alignment in Ten-m null embryos. In contrast to *fas3*^−^ embryos, we observed significant defects in CB arrangements in *ten-m*^−^ embryos, even during CBs migration stages, with the 4-2 pattern of Tin- and Svp-positive CBs perturbed (n=65/87 in *ten-m*^−^, n=16/59 in *fas3*^−^, Figure S6D-S6F”). This indicates the early expression of Ten-m in CBs may have an important role in regulating CBs arrangement, potentially through guiding CBs migration at Stage 13-15 (Raza and Jacobs, 2016) or even affecting progenitor CBs fate determination by ensuring correct division orientation during progenitor CB division at Stage 12 (Tao and Schulz, 2007; Ward and Skeath, 2000). To assess the cell matching defects caused by loss of Ten-m function, we analyzed only mutant embryos with cell arrangement patterns comparable to wildtype. Similar to *fas3*^−^ mutants, loss of Ten-m causes partial reduction in the efficacy of CB matching (Figure 6F-6F’) with the cell mismatch (δ=0.21±0.06) still significantly lower than *svp*^−^ embryos (Figure 6G). Next, we abolished the expression of both Ten-m and Fas3 in double mutant embryos. These embryos showed severe defects in CBs matching with loss of straight boundaries between CB subtypes (Figure 6H-6H’), reminiscent of *svp*^−^ embryos. Quantifying this mismatch, we found CB mismatch is comparable to *svp*^−^ embryos (δ_DM_=0.27±0.10, Figure 6G).

These results support our hypothesis that Fas3 and Ten-m are working complementarily to ensure robust CB matching. Under normal conditions, both Fas3 and Ten-m are regulating cell matching based on their own differential expression pattern (Figure 6I top). The loss of one is compensated by the other, which can still partially function to retain the proper cell alignment (Figure 6I middle). However, when both Fas3 and Ten-m are lost or equally distributed in the heart, CBs dramatically lose the ability to perform active cell matching (Figure 6I bottom), resulting in severe cardiac misalignment.

## Discussion

Precise and robust cell matching is widely required in multicellular organisms and various components have been identified that regulate cell matching (Davenport et al., 1993; Eilken and Adams, 2010; Jacinto et al., 2000; Mossey et al., 2009; Tessier-Lavigne and Goodman, 1996). However, the underlying mechanisms of cell matching, especially with regard to dynamics, remain to be elucidated. Here, we reveal that precise and robust CB matching is mediated by filopodia selective binding activity in distinct cell types and the related complementary differential expressions of the cell-cell adhesion molecules Fas3 and Ten-m (Figure 7). As both Fas3 and Ten-m promote homophilic adhesion (Chiba et al., 1995; Hong et al., 2012; Kose et al., 1997; Mosca, 2015; Snow et al., 1989), we propose that Fas3 and Ten-m instruct robust CB matching through regulating filopodia binding activities, potentially in a competition manner: in wildtype, Tin-positive CBs with high expression of Fas3 lead the heart matching and form stronger filopodia binding with their partner cells, while their filopodia can also reach the matching regions between Svp-positive CBs and disrupt those CBs’ filopodia adhesion, thus creating a differential adhesion affinity and instructing precise matching (Figure 7A); however, in Hand>fas3^RNAi^ embryos, where Fas3 in CBs are significantly reduced, the Svp-positive CBs expressing relatively higher Ten-m become the leader cells and their filopodia are able to form relatively stronger adhesions without the interference from the Tin-positive CBs that lag behind, thus retaining the partial cell matching (Figure 7B).

**Figure 7.**
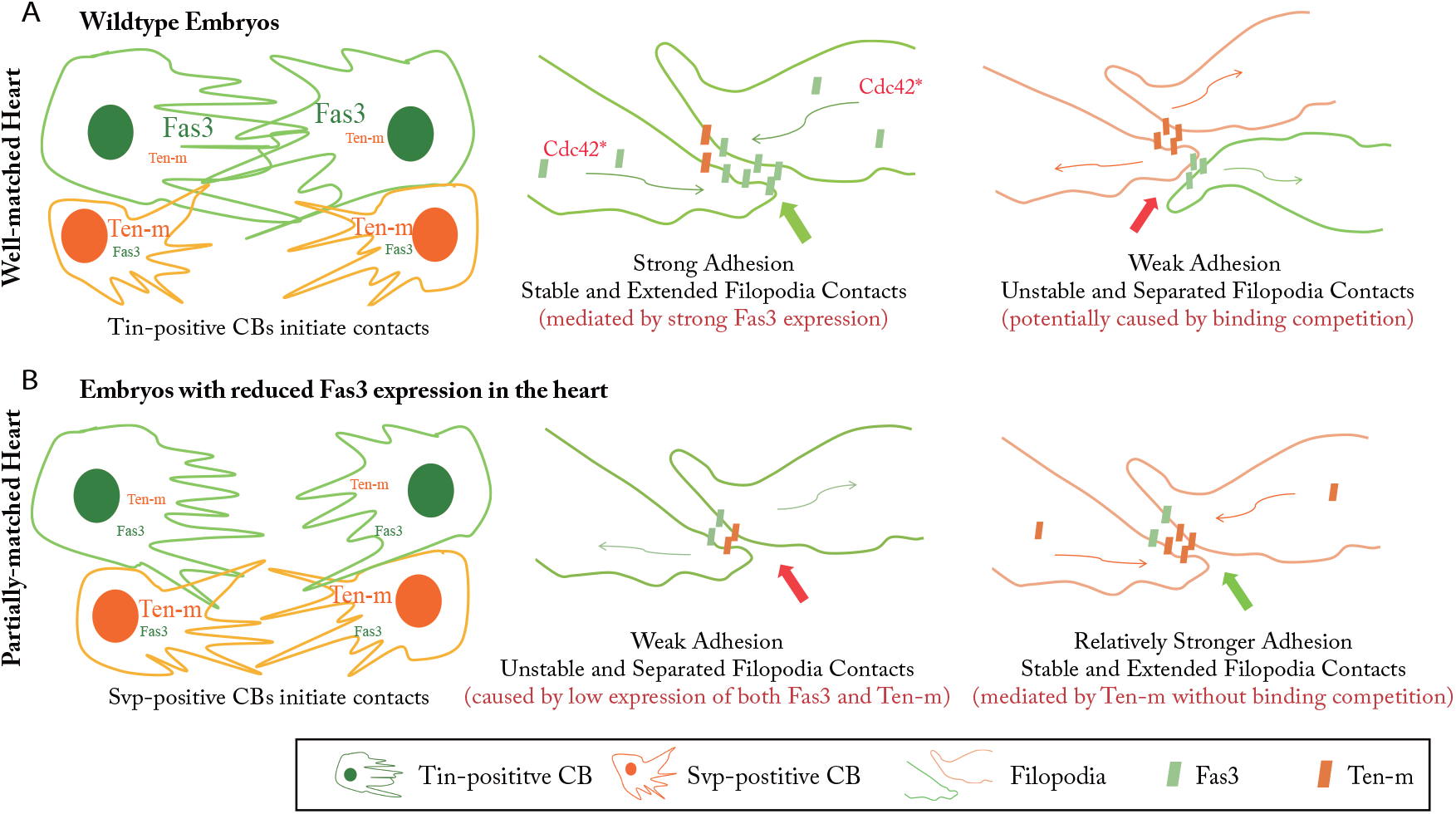
Model of *Drosophila* CB matching through selective filopodia adhesion. (A, B) Schematics of Fas3 and Ten-m mediated CB matching through affecting filopodia binding activities in wildtype(A) and embryos with reduced Fas3 expression in CBs (B). Font sizes of ‘Fas3’ and ‘Ten-m’ indicate the expression levels.

During development, filopodia activity regulates proper cell matching of multiple processes (Davenport et al., 1993; Eilken and Adams, 2010; Jacinto et al., 2000; Tessier-Lavigne and Goodman, 1996). In particular, similar selective filopodia stabilization of developing synapses has also been shown to be important for specific circuit formation (Changeux and Danchin, 1976; Özel et al., 2015; Trachtenberg et al., 2002). Furthermore, the differential expression of cell adhesive molecules has been widely used to explain the formation of cell-cell connections in many tissues, including cell sorting (Amack and Manning, 2012; Fraser, 1980; Gilbert, 2007; Schwabe et al., 2014; Steinberg, 1963). Yet, the link between filopodia activity and differential adhesion - and the subsequent dynamic regulation of cell matching - remains unclear. Here, we have shown that these two components are tightly linked to instruct cell matching and organization; cell specific differential adhesion controls the selective binding of filopodia and thus drives the cells to their correct partners and helps in forming precise cell-cell connections. This provides a simple and efficient matching mechanism that could be generally applicable. For example, in angiogenesis new blood vessel formation involves initial sprouting and rebuilding of connections between blood vessel cells (Adams and Alitalo, 2007); yet, how these cells form precise cell-cell connections remains poorly understood. We propose that blood vessel cells may utilize filopodia selective adhesion to sense the surrounding cellular environment and specifically adhere with their partner cells based on the differential expression of certain cell-cell adhesive molecules.

Functional redundancy is a fundamental principle in building robust biological systems (Hiroaki Kitano, 2004; Masel and Siegal, 2009; Turrigiano, 1999). Here we identify Fas3 and Ten-m as regulating CBs matching in a complementary fashion. Their spatial and temporal expression patterns in CBs appear to be precisely regulated and the degree of adhesion difference is carefully balanced. During neurogenesis, numerous molecules have been identified in guiding cell targeting (Shen and Cowan, 2010; Tessier-Lavigne and Goodman, 1996), and potential complementarity has been demonstrated (Winberg et al., 1998). In other multicellular systems, like neural crest formation (Sauka-Spengler and Bronner-Fraser, 2008), the expression of multiple selective adhesive molecules in different sub-locations is common, and their spatial-temporal regulation is precisely mediated (Gilbert, 2007; Takeichi, 1988; Williams and Barclay, 1988). Combined, we propose that precisely regulated differential adhesion and matching redundancy is an efficient mechanism for ensuring robust cell-cell connection formation.

Although we have revealed how active *Drosophila* CB matching is driven by selective filopodia adhesion, there are a number of open questions. First, how exactly does Fas3 regulate the filopodia binding activity? Is it a simple mechanical process based on homophilic binding, or are further downstream interactions, in particular cytoskeleton remodeling and active adhesion molecule transportation, also involved? Fas3 is likely containing four N-linked glycocylation sites and two phosphorylation sites (Snow et al., 1989), which potentially enables complex regulation after initial filopodia contact. Cdc42 is presumably involved in such processes, but exactly how is still unclear. Second, what is the regulation mechanism of Ten-m and how are these different interactions between filopodia with Fas3 and Ten-m tuned to ensure robust CB matching? Ten-m belongs to the Teneurins, which are conserved throughout Animalia and are related to neuron structures, functions and diseases (Antinucci et al., 2013; Hong et al., 2012; Sklar et al., 2011; Tucker and Chiquet-Ehrismann, 2006). Ten-m facilitates neuron recognition through differentiating between simultaneous homo- and heterophilic interactions (Hong et al., 2012; Mosca, 2015). The downstream effectors of Ten-m and its relationship with Fas3 constitute interesting avenues for further investigations. We believe that future studies on the CB matching process of *Drosophila* heart will shed light on these common challenges in cell matching.

## Author Contributions

S.Z. and T.E.S. designed the project. S.Z. performed the experiments with assistance and advices from C.A.. S.Z. performed the data quantification. All authors analyzed and interpreted the data. All authors contributed to writing the manuscript.

## Acknowledgments

We thank Manfred Frasch, Rolf Bodmer, Stefan Baumgartner, Zhe Han and Yusuke Toyama for sharing fly lines and reagents. We acknowledge Paul Matsudaira, Yusuke Toyama, Ronen Zaidel-Bar, Zhe Han, Adrian Moore and all Saunders’ lab members for fruitful discussions and comments on the manuscript. This work was supported by the National Research Foundation Singapore under an NRF Fellowship to T.E.S. (NRF2012NRF-NRFF001-094).

## Supplemental Information

### Supplementary Figures

**Figure S1.**
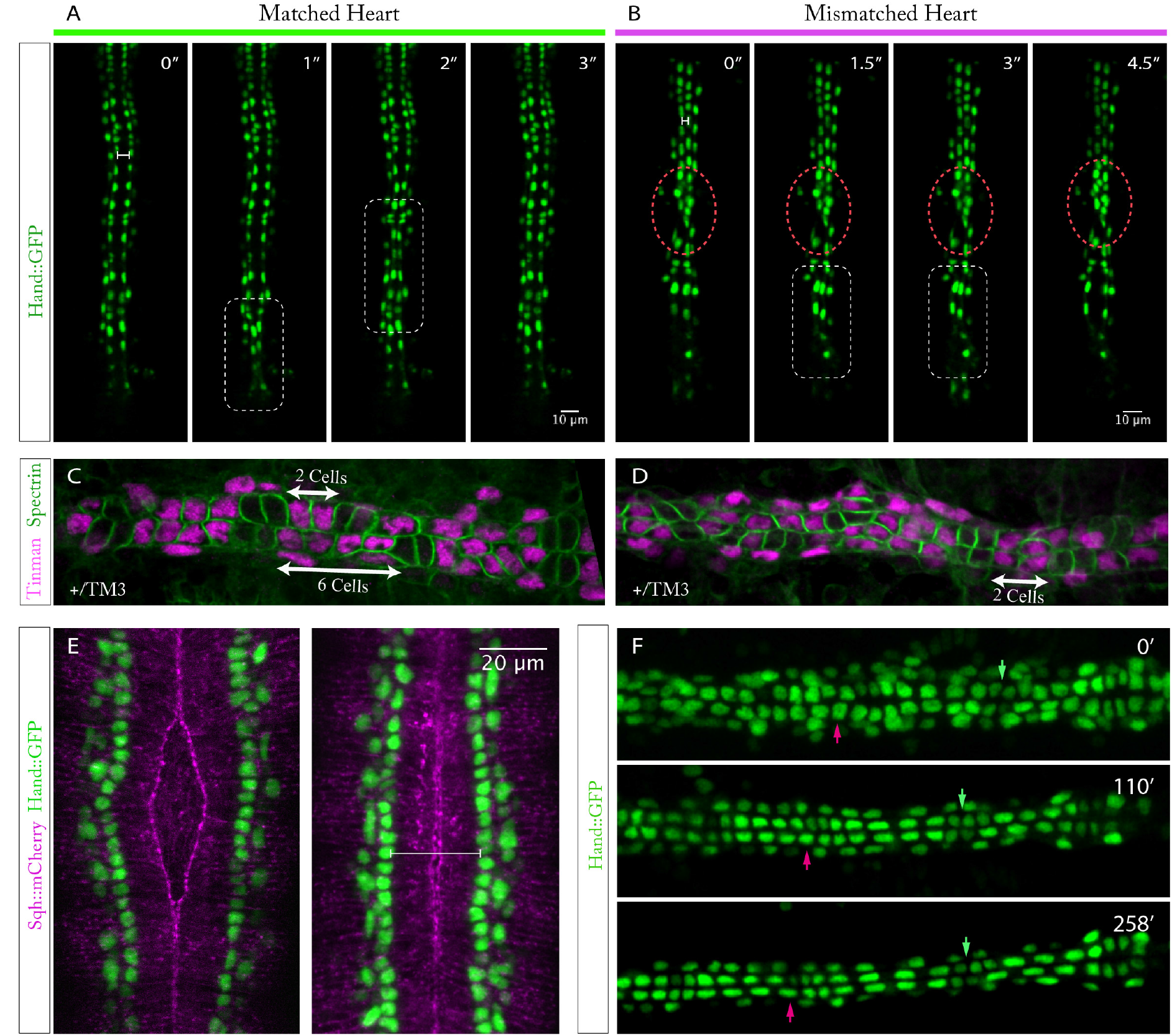
*Drosophila* heart formation is a cell-type specific active matching process, happening in a short time window, related to Figure 1; Movies S1 and S2. (A) Morphology and beating of well aligned heart in Hand::GFP embryos at late Stage 17. (B) Morphology and beating of misaligned heart in Hand::GFP embryos at late Stage 17 (n=5). (A-B) Red circles label the misaligned CBs regions, white boxes label the heart contraction regions, white bars label the contralateral CBs separation during the relax state. (C-D) CBs arrangement and alignment in +/TM3 embryos. (E) Relative position of CBs (Hand::GFP labeled) migration and epidermis dorsal closure (Sqh::mCherry labeled). White bar labels the contralateral CB separation at the time of dorsal closure finishing. (F) CBs morphology and alignment after full coalescence of contralateral CBs in Hand::GFP embryos. Green and red arrows point to the matched and mismatched CBs.

**Figure S2.**
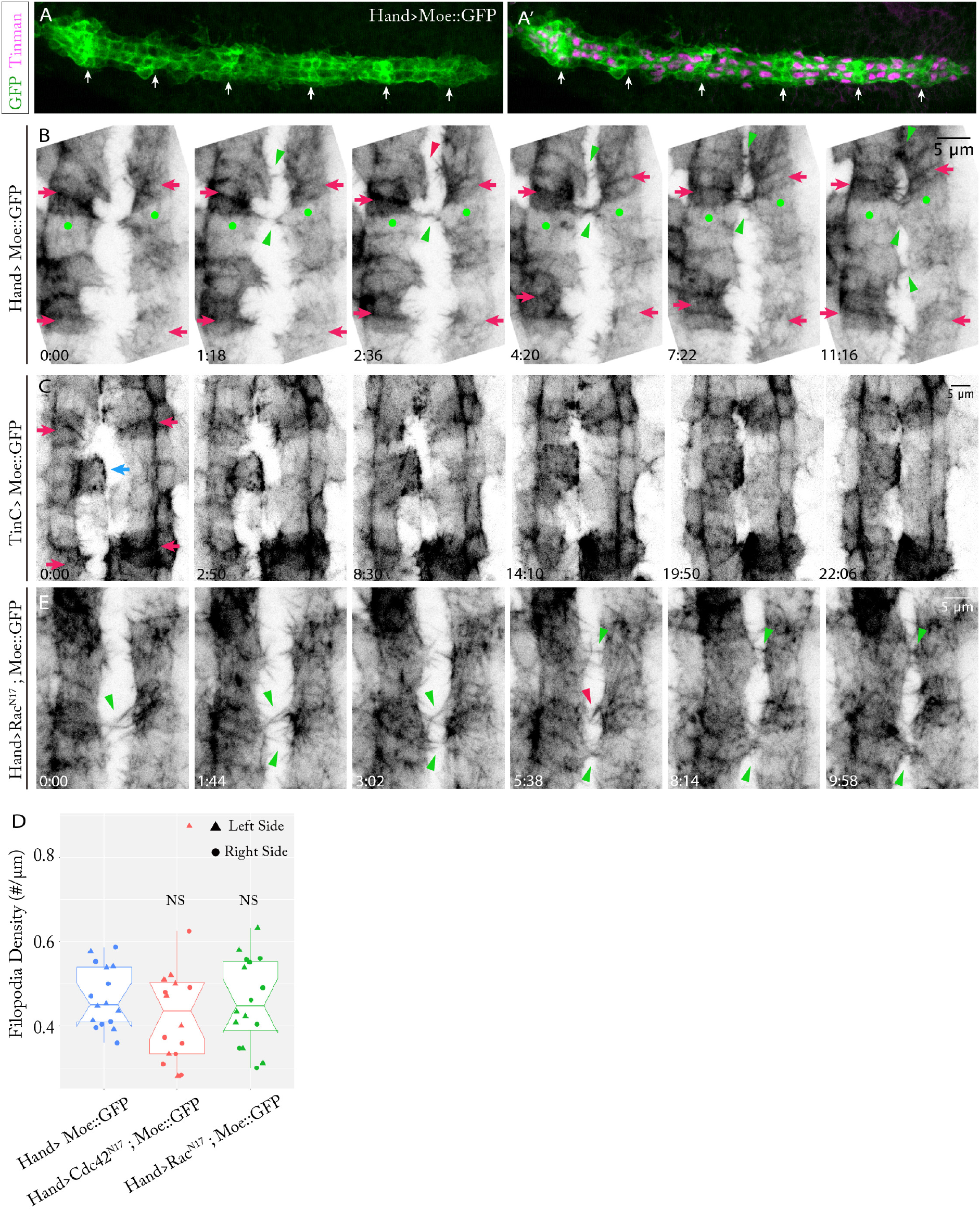
CB filopodia show selective binding affinity, related to Figure 2. (A-A’) Expression pattern of Moe::GFP in CBs when driven by Hand-Gal4. White arrows point to the Svp-positive CBs. (B) Filopodia binding activity in an Hand>Moe::GFP embryo where different CB subtypes can be clearly distinguished. Red arrows point to the distinguished Svp-positive CBs, green dots point to the Tin-positive CBs that makes strong filopodia binding contacts. (C) Moe::GFP expression in TinC>Moe::GFP embryo. Red and blue arrows point to the distinguished Svp-positive CBs and the surrounding aminoserosa cells. (D) Filopodia density quantification in different conditions. Data are presented as boxplot and scatter plot (n=8 per condition) (NS: Not Significant, compared with Ctrl condition). (E) Filopodia binding activity in Hand>Rac^N17^; Moe::GFP embryo. (B, E) Green and red arrowheads point to the established and separated filopodia contacts.

**Figure S3.**
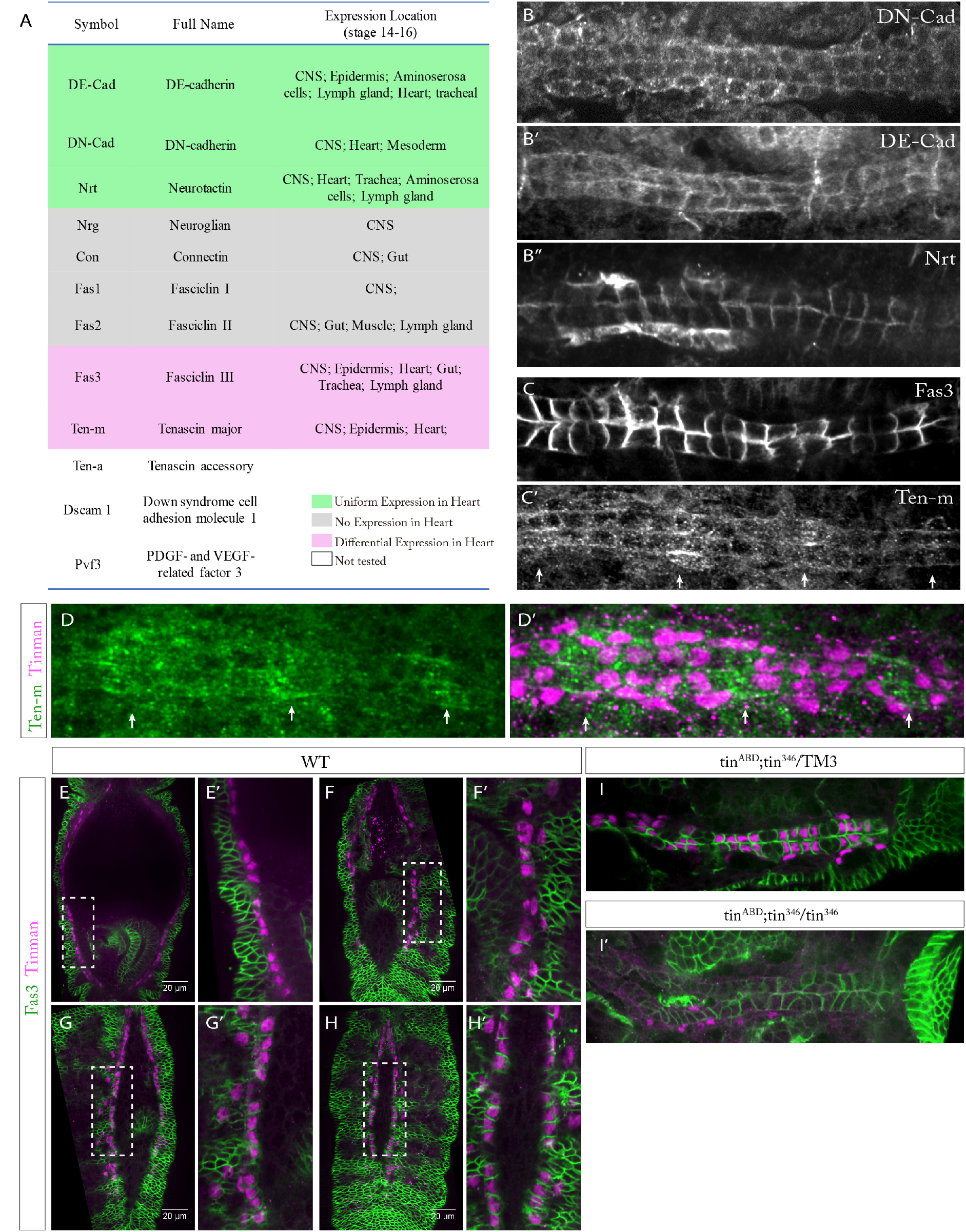
Candidate screening identified differential expression of Fas3 and Ten-m in CBs, related to Figure 3. (A) List of candidate matching molecules and their expression locations in the embryos at stage 14-16. CNS: Central Nervous System. (B-B”’) Expression pattern of candidate molecules DE-Cad (B), DN-Cad (B’) and Nrt (B”) in CBs of wild-type embryos at Stage 16. (C, C’) Expression pattern of Fas3 (C) and Ten-m (C’) in CBs of wild-type embryos at stage 16. (D, D’) Ten-m and Tin staining in wild-type embryos at Stage 16. (E-H’) Fas3 expression in the CBs at Stage 14 (E, E’), early Stage 15 (F, F’), middle Stage 15 (G, G’) and late Stage 15 (H, H’). (E’-H’) Magnified images of the white line labeled regions in (D-G). (I, I’) Fas3 expression pattern and CB morphology in heterozygous (I) and homozygous (I’) *tin* mutant embryos.

**Figure S4.**
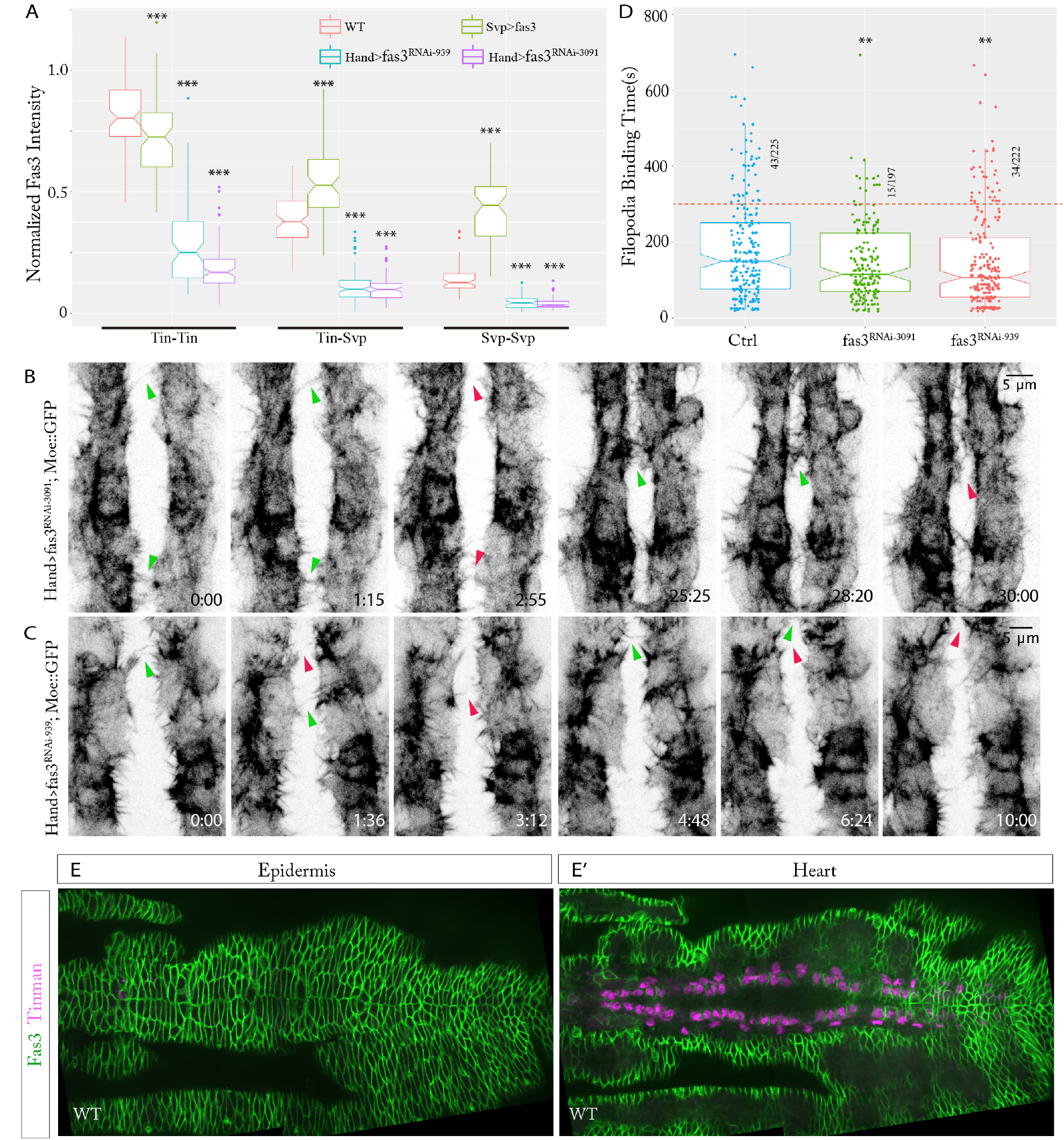
fas3^RNAi^ efficiently decreases the Fas3 expression in CBs and reduces CB filopodia binding affinity, related to Figure 3 and Movie S3-S4. (A) Fas3 intensity quantification at different cell-cell contacts in different experimental conditions (n=7 embryos per condition). Data are presented as boxplot (*P<0.05, **P<0.01, ***P<0.001 compared to the wildtype at each different cell-cell contact). (B, C) Filopodia activities in Hand>fas3^RNAi-3091^ (B) and Hand>fas3^RNAi-939^ (C) embryos. Green and red arrowheads point to the established filopodia bindings and the separated ones. (D) Filopodia binding time in different conditions. Data are presented as boxplot and scatter plot (*P<0.05, **P<0.01, ***P<0.001), red line shows the 300s criterion for strong binding. (E, E’) Fas3 expression in *Drosophila* epidermis (E) and heart (E’) of the same embryo.

**Figure S5.**
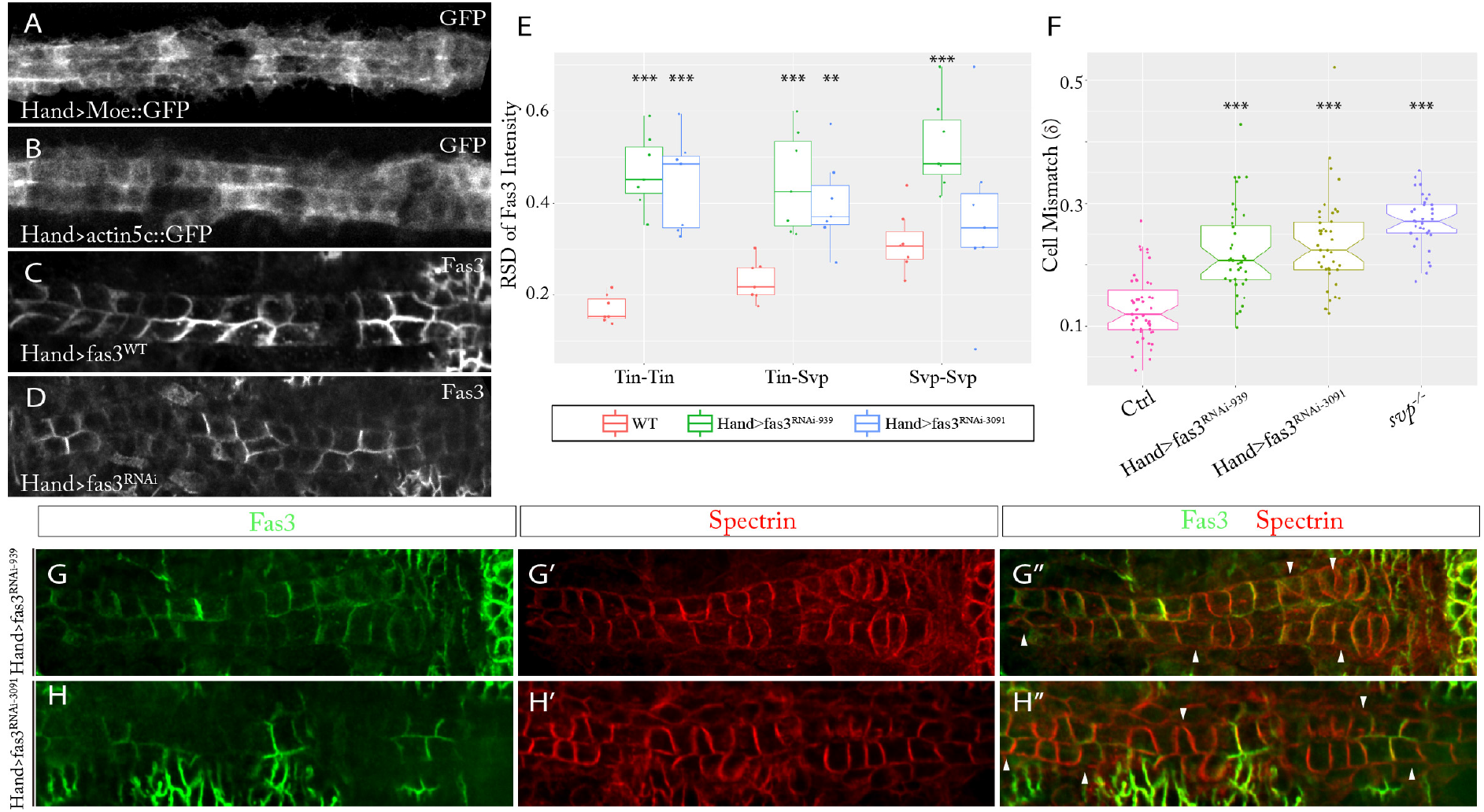
Hand>Gal4 shows stochastic activity and perturbation of Fas3 expression pattern alters CBs matching, related to Figure 3 and Figure 4. (A-C) Expression patterns of Moe::GFP (A), act5c::GFP (B) and Fas3 (C) in the heart driven by the same Hand-Gal4. (D) fas3^RNAi^ driven by Hand-Gal4 generates a stochastic Fas3 expression pattern in the heart. (E) Relative standard deviation of normalized Fas3 intensity at different cell contacts in different experimental conditions (n=7 per condition). (F) CBs Mismatch (δ) analysis of different experimental conditions (n>20 per condition). (E, F) Data are presented as boxplot and scatterplot (*P<0.05, **P<0.01, ***P<0.001 compared to the wildtype). (G-H”) Fas3 expression pattern and CBs alignment in Hand>fas3^RNAi-939^ (G-G”) and Hand>fas3^RNAi-^3091 (H-H”) embryos with different fas3^RNAi^ driven by Hand-Gal4. White arrowheads point to the cells with obvious mismatch.

**Figure S6.**
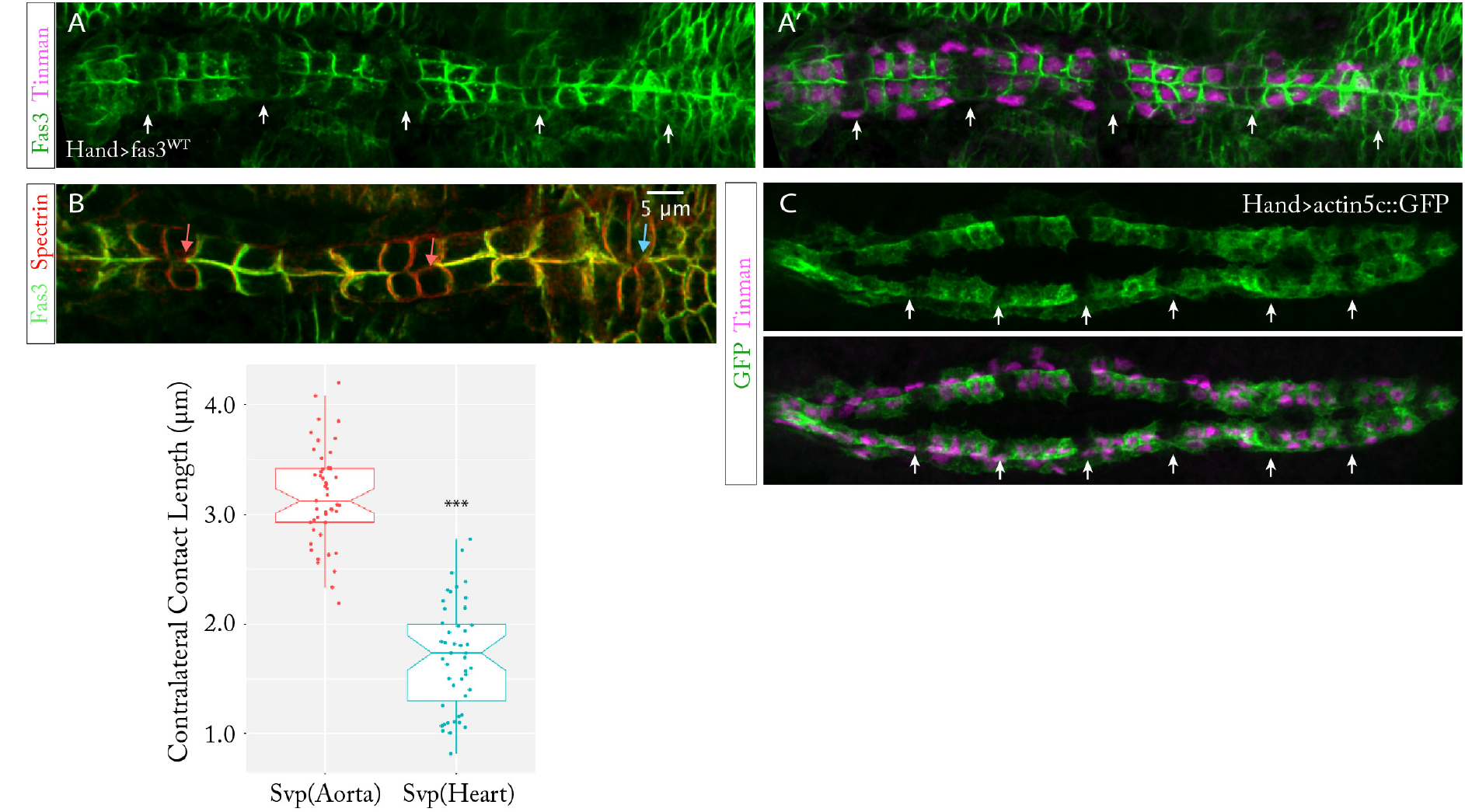
Low expression of Fas3 in Svp-positive CBs is essential for robust heart morphology, related to Figure 5. (A, A’) Fas3 expression in CBs with Fas3 overexpression driven by Hand-Gal4. (B) Contralateral contact length of Svp-positive CBs at the aorta (red arrows pointed) and heart (blue arrow pointed) domains. Data are presented as boxplot and scatterplot (*P<0.05, **P<0.01, ***P<0.001). (C) Act5c::GFP expression in the CBs when driven by Hand>Gal4. (A, A’, C) White arrows point to the locations of Svp-positive CBs.

**Figure S7.**
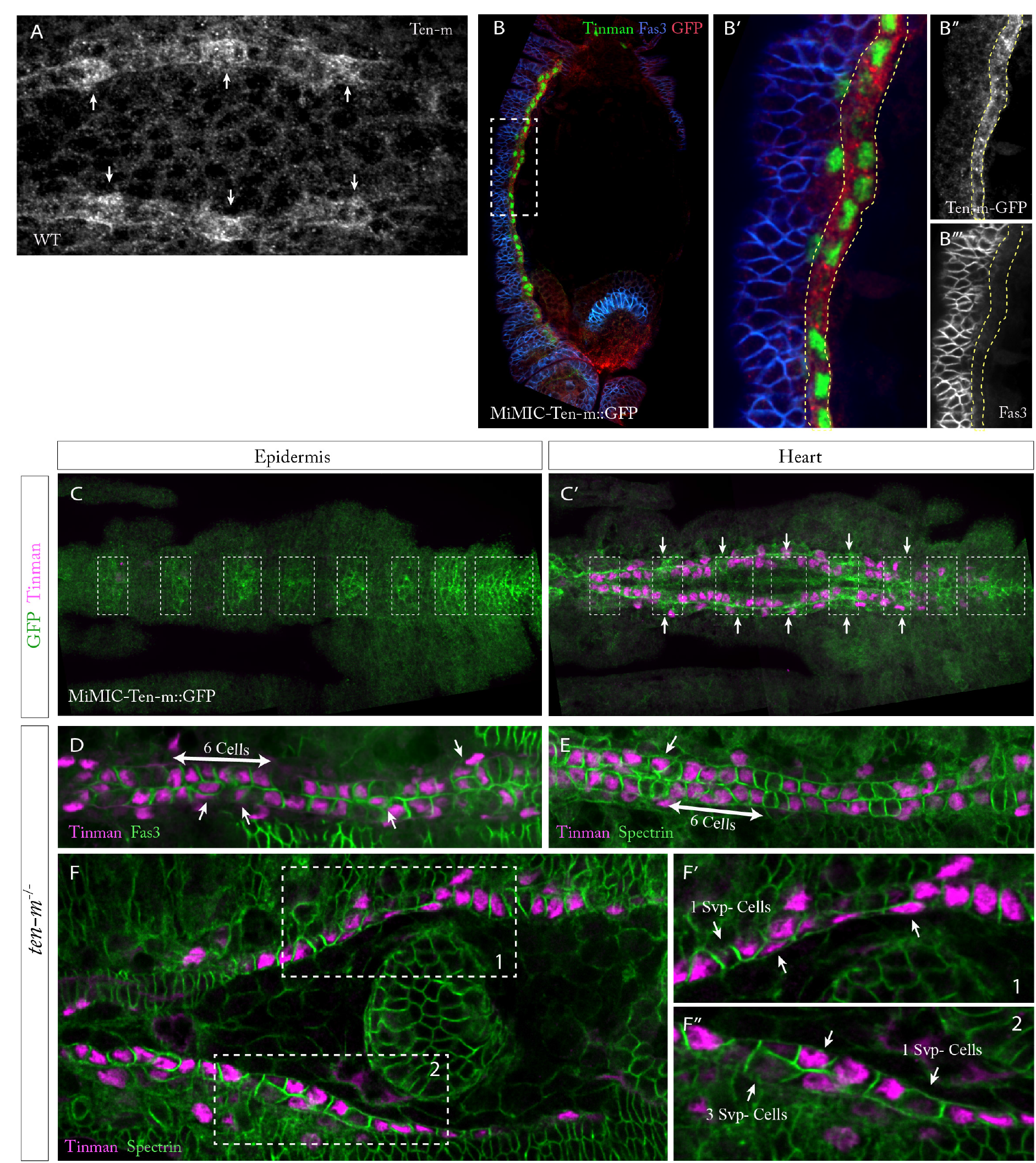
Ten-m shows differential expression in the heart and loss of Ten-m causes CBs arrangement defects, related to Figure 6. (A) Ten-m staining image of wild-type embryo heart at Stage 15. White arrows point to the high Ten-m expression regions. (B) Expression of Ten-m and Fas3 in the heart at the beginning of Stage 14. (B’) Magnified image of the white box labelled region in B. (B”) Ten-m expression channel in (B’). (B”’) Fas3 expression channel in (B’). (B’-B”’) Yellow lines label out the regions of CBs location. (C, C’) Localized expression of Ten-m in epidermis (C) and heart (C’) in the same embryo. White boxes label the high Ten-m expression regions in epidermis, white arrows point to the Svp-positive CBs locations. (D-F) CBs arrangement defect in ten-m mutant embryos at Stage 16 and Stage 15. (F’-F”) Magnified images of the white boxes labelled regions in G. White arrows point to the regions with CBs miss-arrangement.

## STAR Methods

### Fly stocks and genetics

Fly stocks were maintained at 25°C on standard fly food. The following fly stocks were used: Sqh::mCherry (a gift from Yusuke Toyama’s Lab). Hand::GFP, Hand-Gal4 (R25) (a gift from Zhe Han’s Lab), tin^ABD^; Tin^346^/TM3 (a gift from Manfred Frasch’s Lab), TinC>Gal4 (a gift from Rolf Bodmer’s Lab). Fly lines from the Bloomington Drosophila Stock Center: Svp>Gal4 (BL #47912), Svp[AE127] (BL #26669), UAS>Moe::GFP (BL #31776), Fas3[E25] (BL #4748), Mimic-Ten-m::GFP (BL #59798), Ten-m [5309] (BL #11657), UAS>Cdc42^N17^ (BL #6288), UAS>Rac^N17^ (BL #6292), UAS>act5c::GFP (BL #9257), TM6b-Ubi::GFP (from BL #4887, used for distinguishing homozygous Ten-m mutant). Both RNAi lines targeting *fas3* from the Vienna *Drosophila* RNAi Center: UAS>fas3^RNAi^ (VDRC ID3091, ID939).

Hand::GFP is a transgene carrying a GFP reporter driven by the Hand cardiac and hematopoietic (HCH) enhancer (Han, 2006). UAS-Moe::GFP is a transgene with GFP tagged to the actin-binding domain of moesin (Dutta et al., 2002).

A UAS-Fas3 fly line was generated by P-element transformation, carried out by BestGene Inc. The UASp-Fas3 plasmid was a a gift from Victor Hatini (de Madrid et al., 2015).

### Stochastic activity of Hand-Gal4 in the *Drosophila* heart

Gene expression (Moe::GFP, act5c::GFP, fas3 tested here) driven Hand>Gal4 results in stochastic expression pattern, especially among the cells of same CB subtype (Figure S5A-S5D). Similarly, this variable and unstable Gal4 activity has also been very recently described in other enhaner-Gal4 lines (Casas-Tintó et al., 2017). Likely, this stochastic Gal4 activity is related to the primary sequence of insertion site as well as the complex three-dimensional structures of the chromosome. Moreover, when driven by Hand>Moe::GFP shows higher expression in Svp-positive CBs (Figures S2A and S2A’), but act5c::GFP (Figure S5K) and Fas3 (Figures S5I and S5I’) show higher expression in Tin-positive CBs. This indicates that post-transcriptional regulation potentially exists for these different molecules in distinct CB subtypes.

### Immunostaining

*Drosophila* embryos were collected at the desired stage, dechorionated in household bleach and fixed in 4% formaldehyde and blocked with 10% BSA in PBT (0.1% Triton in PBS) according to standard procedures. To do co-staining of Fas3 and Spectrin (both were generated in mouse host), sequential staining was performed – after the staining of Fas3 using the standard procedures, 4% PFA (in PBS) was used to post-fix the embryos for 20mins, then staining of Spectrin was continued in standard methods. The following primary antibodies were used in this study: rabbit anti-Tinman (1:1000), chicken anti-GFP (1:1000, ThermoFisher Cat#A10262/ invected RRID: AB_2534023), mouse anti-Spectrin (1:400, DSHB Cat# 3A9 (323 or M10-2) /invected RRID: AB_528473), rat anti-ECad (1:300, DSHB Cat# DCAD2/invected RRID: AB_528120), rat anti-CadN (1:300, DSHB Cat# DN-Ex #8/invected RRID: AB_528121), mouse anti-Nrg (1:300, DSHB Cat# BP 104 anti-Neuroglian /invected RRID: AB_528402), mouse anti-Nrt (1:300, DSHB Cat# BP 106 anti-Neurotactin /invected RRID: AB_528404), mouse anti-Con (1:300, DSHB Cat# Connectin C1.427 /invected RRID: AB_10660830), mouse anti-Fas1 (1:30, DSHB Cat# F5H7 anti-Fasciclin I /invected RRID: AB_528233), mouse anti-Fas2 (1:300, DSHB Cat# 1D4 anti-Fasciclin II/invected RRID: AB_528235), mouse anti-Fas3 (1:300, DSHB Cat# 7G10 anti-Fasciclin III/invected RRID: AB_528238, monoclonal anti-Ten-m (Mab113, 1:100). Primary antibodies were detected with Alexa Fluor-labelled secondary antibodies (1:500; LifeTech). Embryos were imaged on a Nikon SpinningDisk_W1_LiveSR with a CFI Plan Apochromat 100x/1.45 NA oil immersion objective. Tinman antibody was kindly provided by Manfred Frasch. Ten-m antibody was a a gift from Stefan Baumgartner.

### Live Imaging

Embryos were collected, dechorionated, mounted on a MatTek dish and imaged on Zeiss LSM710 microscope. For the imaging of Hand::GFP; histone::mCherry lines, embryos were imaged using a C-Apochromat 40x/1.2 NA water-immersion objective with an interval time of 5mins. Hand::GFP labelled heart beating was imaged using a C-Apochromat 40x/1.2 NA water-immersion objective with interval time 1s. Hand::GFP; sqh>Sqh::mCherry embryos were imaged using a C-Apochromat 63x/1.2 NA water-immersion objective with interval time 2mins. All UAS-Moe::GFP labelled filopioda activities were imaged using C-Apochromat 63x/1.2 NA water-immersion objective with interval time 20-30 seconds.

### Cell tracking

Tracking of Hand::GFP labelled CBs migration over long time was done in a custom-built software in Matlab R2015a. Before analysis, each Z-stack for each time point was Z-projected (maximum intensity) in ImageJ. In the tracking software, individual CB at each time point was detected using the ‘imfindcircles’ function in Matlab, and the position information (including x, y, t) of each cell centre (P^t^) stored. The image at frame t+1 was compared to frame t and analysed with Particle Image Velocimetry (PIV) algorithm (original code from ‘OpenPIV’)(Taylor et al., 2010). The predicted new position (P^p^) of each cell centre (P^t^) at frame t+1 (P^t+1^) was derived based on the predicted velocity results from PIV analysis. To decrease the prediction noise in PIV, the averaged predicted velocity of a 3×3 point window (with P^t^ centred) was used for the predicted velocity (P^v^). The predicted cell centre position at frame t+1 (P^p^)=P^t^+P^v^. Next the tracked position of P^t^ at frame t+1 (P^t+1^) was assigned to the nearest detected cell centre at frame t+1 to P^p^. Manual correction was performed to further increase the tracking precision.

Individual cell tracking shown in SF.2 C” was done in Bitplane Imaris 8.2.0. Images were segmented using automatic spots detection for GFP channel and adjusting threshold accordingly. Cell tracking was performed by using the autoregressive motion algorithm.

### Mismatch Quantification

Mismatch measurement based on membrane contacts was done in ImageJ. Partner cells were assigned by their cell type marker and relative position in the repeated 4-2 cell arrangement. In *svp*^−^ mutant, the cell from contralateral side that formed the largest contact was assigned as the partner of the cell being measured. Junction points between cell lateral boundaries and the middle contact line of the two contralateral sides were manually labeled out. Each contact length between two neighbored points was estimated by their direct distance. Length of these contacts was assigned to different cells and classified into ‘matched contact’ or ‘mismatched contact’ based on whether it is between two partner cells or not. Mismatch rate is calculated by dividing the mismatched contact length with total membrane contact length. With this definition, the mismatch value ranged between 0 and 0.5.

Mismatch measurement based on cell tracking was done in Matlab. Through the automatic cell tracking, the cell positions were stored. To increase the measurement quality, only the tracked cells in the middle region (segments ~A2-A5) were used for this analysis. Partner cells were defined in the last time frame, cells from contralateral sides with minimum center distance separation were assigned as partner cells. Midline at each frame was defined as the regression line of center points between each partner cells. Each cell center was projected towards the midline, and the distance between projected lines of two partner cells was assigned as the mismatch distance. The mismatch was then calculated by dividing the averaged mismatch distance by the averaged distance of neighboring cells.

### Quantification of filopodia binding time

Moe::GFP labeled filopodia binding activities were visualized and analyzed using Zen Lite 2012 (Blue edition). The filopodia contact initiation time frame f^in^ and the separation time frame f^end^ were recorded. For the contacts that held until the full cell fusion, the time frame that the filopodia structure is not distinguishable is recorded as f^end^. The filopodia binding time was calculated, binding time=image interval time × (f^end^ - f^in^).

### Quantification of Normalized Fas3 Intensity

Fas3 intensity was measured using ImageJ. Fas3 intensity at epidermis cell-cell boundaries was measured, averaged and defined as I_e_. Fas3 intensity in non-Fas3 expressing cells in the same embryo was measured, averaged and defined as the background intensity, I_b_. Fas3 intensity at CB-CB boundaries (I_CB_) in the same embryo was measured and normalized (I_NCB_) to 0-1 according to its relative value to I_e_ and I_b_: I_NCB_= (I_CB_ - I_b_)/(I_e_ - I_b_).

### Statistical analysis

Statistical analysis was performed with Matlab using two sample t-test. Plots were generated using R. When comparing fraction of embryos displaying phenotypes, the p-value was calculated using a Chi-squared test.

## SUPPLEMENTAL MOVIE TITLES AND LEGENDS

**Movie S1. Heart morphology and beating in Hand::GFP embryos, related to Figure S1**

Time-lapse image of Hand::GFP (green color) labelled heart beating with well aligned (left) and misaligned (right) CBs. Time given in min:sec. The same data are presented as montage in Figures S1A and Figure S1B, respectively.

**Movie S2. Relative position of CBs migration and dorsal closure, related to Figure1 and Figure S1**

(Left) Time-lapse image of Hand::GFP (green color) labelled CBs migration and their relative position with dorsal closure (Sqh::mCherry labelled, magenta color). (Right) Cell tracking of individual CBs using the same movie (the cell traces are labelled with migration traces color-coded by migration speed). The same data are presented as montage in Figures 1I and Figure S1E.

**Movie S3. Filopodia activity in control, fas3^RNAi^, and Cdc42^N17^ expressed CBs, related to Figure 2 and Figure 3**

Time-lapse image of Moe::GFP marked CBs actin cytoskeleton, especially filopoida, activity in Hand>Moe::GFP (Left), Hand>fas^RNAi^; Moe::GFP (middle), Hand>Cdc42^N17^; Moe::GFP (right) embryos. Time given in min:sec. The same data are presented as montage in Figures 2B, 2E and Figure 3F respectively.

**Movie S4. Filopodia activity in control and Fas3 overexpressed Svp-positive CBs, related to Figure 2 and Figure 3**

Time-lapse image of Moe::GFP marked CBs actin cytoskeleton, especially filopodia, activity in Svp>Moe::GFP (Left), Svp>fas3^WT^; Moe::GFP (right) embryos. Time given in min:sec. The same data are presented as montage in Figures 2D and Figure 3I, respectively.

**Movie S5. CBs migration pattern in control and fas3^RNAi^ expressed embryos, related to Figure3**

Time-lapse image of Moe::GFP marked CBs migration in Hand>Moe::GFP (Left), Hand> fas^RNAi^; Moe::GFP (right) embryos. Time given in min:sec. The same data are presented as montage in Figures 3J and Figure 3K, respectively.

**Movie S6. Filopoida activity of Tin-positive CBs neighboring Svp-positive CBs in Hand>act5c::GFP embryos, related to Figure 5**

Time-lapse image of act-5c::GFP marked CBs filopodia activity in Hand>act-5c::GFP embryos. Time given in min:sec. The same data are presented as montage in Figures 5F.

